# *Aeromonas* adhesins facilitate kin and non-kin attachment to enable T6SS-mediated antagonism in liquid

**DOI:** 10.64898/2026.01.27.701733

**Authors:** Chaya Mushka Fridman, Hadar Cohen, Kinga Keppel, Alejandro Tejada-Arranz, Marek Basler, Motti Gerlic, Dor Salomon

## Abstract

Bacterial ability to deploy the type VI secretion system (T6SS) against rivals requires prolonged cell-cell interactions. Such interactions are facilitated on solid surfaces but are assumed to be absent in liquid, leading to the conventional dismissal of T6SS-mediated competition in liquid environments. Here, we find that *Aeromonas jandaei* employs its T6SS to eliminate diverse bacterial competitors in liquid media. Using a workflow that monitors interbacterial competition via prey luminescence, we demonstrate that auto-aggregation and co-aggregation, facilitated by distinct adhesins, enable kin and non-kin recognition and intoxication in a T6SS-dependent manner. Furthermore, we show that another marine bacterium, *Vibrio coralliilyticus*, employs T6SS to intoxicate rivals in liquid media. Collectively, our results indicate that T6SS-mediated competition in liquid is more common in marine bacteria than previously anticipated, and can be facilitated by diverse molecular mechanisms that govern cell aggregation.

## INTRODUCTION

A common weapon used in interbacterial competition is the Type VI secretion system (T6SS), a widespread protein delivery apparatus^1–5^. Most T6SSs studied to date deliver cocktails of antibacterial toxins, called effectors, into neighboring bacteria in a contact-dependent manner^2,6–10^. To this end, a tube-spike structure, decorated with effectors, is propelled out of the cell by the contraction of an engulfing sheath^11^. The tube-spike then penetrates a neighboring bacterium and deploys the effectors. Several T6SSs were shown to deliver anti-eukaryotic effectors into eukaryotic cells, thereby implicating these systems in virulence, defense against predation, or competition with eukaryotic microbes^12–16^.

Many antibacterial T6SS effector families have been investigated, revealing toxic mechanisms and targets in the bacterial cytoplasm, periplasm, and membrane, including peptidoglycan-degrading enzymes^6,17,18^, poreforming toxins^19–24^, phospholipases^8,25^, nucleases^26–29^, NAD(P)^+^-degrading enzymes^30,31^, ADP-ribosyl transferases^32,33^, and effectors inhibiting protein synthesis^34^, DNA replication^35^, or cell division^32^. Importantly, antibacterial effectors are encoded adjacent to a cognate immunity protein that prevents self- or kin-intoxication^2,36–38^. Nevertheless, several non-immunity protein-mediated defense mechanisms that nullify or attenuate T6SS-mediated toxicity have been recently revealed^34,39–49^, and it is likely that other such mechanisms exist^46,50^.

Until recently, it had been widely accepted that T6SS-mediated interbacterial competition does not occur in liquid environments since it requires prolonged cell-cell contact, which is forced on solid surfaces^51,52^. However, three recent reports challenge this conviction: (1) *Vibrio fischeri* was shown to aggregate in liquid media containing calcium, thus enabling T6SS-mediated intra-species competition^53^; (2) *Vibrio cholerae* was shown to use type IV pili to facilitate cell-cell attachment in liquid, thus enabling T6SS-mediated intra-species competition^54^; (3) the aquatic filamentous bacterium *Aureispira* sp. CCB-QB1 was proposed to use a type IX secretion system (T9SS) to grapple onto non-kin prey bacteria, facilitating T6SS-dependent predation in liquid^55^. These findings raise an intriguing question: Can other bacteria use the T6SS to compete in liquid environments? If so, what conditions enable T6SS-mediated killing in liquid, and what mechanisms are used to facilitate prolonged cell-cell interactions? Furthermore, the aforementioned studies observed either inter- or intra-species T6SS-mediated competition in liquid; it remains unknown whether attacking cells distinguish between kin and non-kin prey cells for attachment in liquid. Answers to these questions may revise current concepts of the roles played by T6SS from evolutionary and ecological perspectives, especially in aquatic bacteria, many of which are known or emerging human and animal pathogens^56,57^.

Here, we report that the bacterium *Aeromonas jandaei* uses specific adhesins to attach to kin or non-kin prey cells in liquid media and intoxicate them in a T6SS-dependent manner. Furthermore, we identify additional marine bacteria that mediate T6SS-dependent interbacterial competition in liquid media, thereby revealing a common phenomenon in aquatic bacteria.

## RESULTS

### Aeromonas jandaei uses T6SS to kill kin prey in liquid media

In a recent study, we found that the T6SS encoded by *Aeromonas jandaei* DSM 7311 (*Aj*) is an antibacterial weapon deploying at least four effectors to antagonize rivals^58^. While investigating the effector Awe1, a peptidoglycan-degrading enzyme, we made an unexpected observation. We initially set out to monitor Awe1 secretion when expressed from an inducible plasmid in a strain in which the endogenous effector and its cognate immunity genes have been deleted (*Aj*Δ*awei1*). We reasoned that since Awe1 targets the bacterial periplasm, its expression in the cytoplasm of a secreting cell should not be toxic even in the absence of the immunity proteins; moreover, we reasoned that it should not intoxicate the neighboring sensitive kin bacteria, since it is widely acknowledged that T6SS does not mediate competition during growth in liquid media due to insufficient cell-cell interactions^51^. Unexpectedly, when monitoring bacterial growth in liquid media, we observed Awe1-mediated toxicity in the absence of its cognate immunity proteins (i.e., in *Aj*Δ*awei1*).

Notably, this toxicity was dependent on a functional T6SS, as inactivation of the T6SS by deleting the sheath component *tssB* restored normal growth (Fig. 1A and Supplementary Fig. S1). This result led us to hypothesize that, contrary to the widely accepted notion^51^, *Aj* uses its T6SS to mediate interbacterial competition in liquid media.

**Figure 1.**
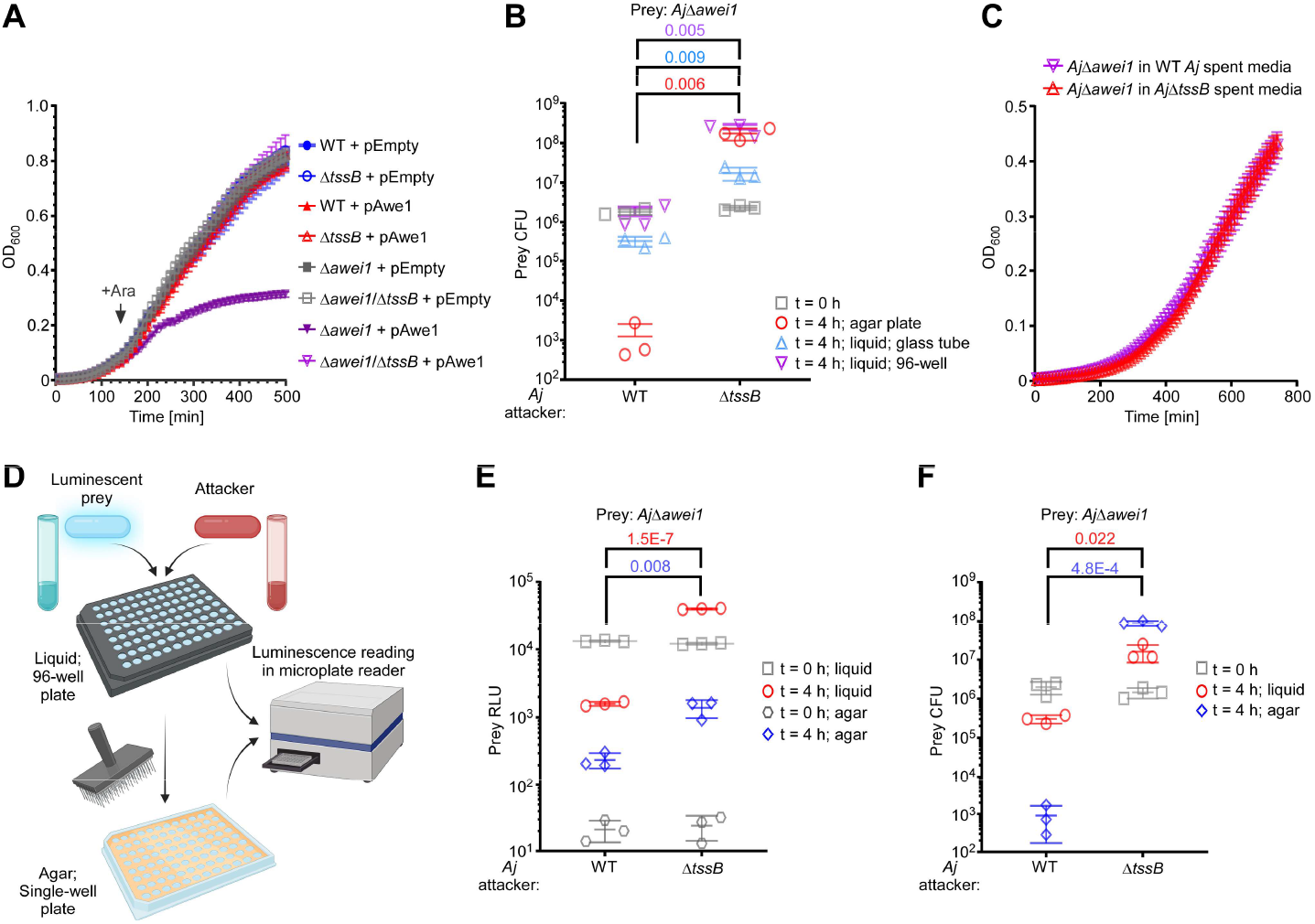
*Aeromonas jandaei* employs its T6SS to intoxicate kin competitors in liquid media. **(A)** Growth of the indicated *Aj* strains containing an arabinose-inducible expression plasmid, either empty (pEmpty) or encoding Awe1 (pAwe1), in LB media at 30^°^C. An arrow denotes the time point at which arabinose (0.05% [w/v]) was added to the media. **(B)** Viability counts (colony forming units; CFU) of *Aj*Δ*awei1* prey before (t = 0 h) and after (t = 4 h) co-incubation with the indicated *Aj* attacker at a 4:1 (attacker:prey) ratio. **(C)** Growth of *Aj*Δ*awei1* in a 1:1 mixture of LB media and the indicated spent media at 30^°^C. **(D)** Illustration of the LiQuoR workflow. **(E)** LiQuoR-based viability measurements (relative luminescence units; RLU) of *Aj*Δ*awei1* prey before (t = 0 h) and after (t = 4 h) co-incubation with the indicated *Aj* attacker at a 4:1 (attacker:prey) ratio. **(F)** Same as E, except prey viability was determined by CFU counts. Data are shown as the mean ± SD; n = 4 (A) or n = 3 (B-C, E-F) samples. The statistical significance between samples at the 4 h time point (colored according to the treatment) was calculated using an unpaired, two-tailed Student’s *t* test. The experiments were repeated three times with similar results. Results from a representative experiment are shown. WT, wild-type.

To directly examine our hypothesis, we monitored the viability (as colony-forming units; CFU) of *Aj* prey that is sensitive to Awe1-mediated toxicity (*Aj*Δ*awei1*) during co-incubation with *Aj* attackers, either wild-type or T6SS^−^ (Δ*tssB*), in a glass tube containing liquid media (Lysogeny broth; LB) at 30^°^C with shaking. Indeed, we observed T6SS-dependent intoxication of the Awe1-sensitive prey after four hours of co-incubation with parental *Aj* attackers (Fig. 1B), confirming that *Aj* uses T6SS to mediate interbacterial competition in liquid. Furthermore, we found that competition in liquid media also occurs when bacteria are incubated in 96-well plates containing LB media, with shaking (Fig. 1B). Therefore, we decided to perform all subsequent liquid media competition assays in 96-well plates.

Next, we sought to determine whether the observed T6SS-mediated competition in liquid is contact-dependent, as an alternative explanation would be that secreted T6SS effectors can enter prey cells independently of direct injection by the T6SS apparatus. To test the latter possibility, we examined whether media containing *Aj* T6SS effectors can impair the growth of an Awe1-sensitive prey strain. To this end, we monitored the growth of the *Aj*Δ*awei1* strain in cleared spent media in which either wild-type or T6SS^−^ *Aj* strains were grown for four hours. As shown in Fig. 1C, the presence of T6SS effectors in the wild-type *Aj* spent media had no effect on the growth of *Aj*Δ*awei1* prey, indicating that the observed T6SS-mediated competition in liquid is contact-dependent.

### Prey viability during competition can be evaluated using luminescence

Evaluating prey viability using CFU counts is labor-intensive. Furthermore, a recent report suggested that estimating T6SS activity using CFU counts on selective media containing antibiotics may lead to an overestimation of the antagonistic effect^59^. Therefore, we sought an alternative reporter that would enable real-time quantification of prey viability during competition in both liquid and solid environments. We decided to use luminescence, which is based on an enzymatic reaction, as an indicator of prey viability that is easy to monitor and quantify^60^. To this end, we introduced a plasmid constitutively expressing the bacterial *Lux* operon^60^ into our prey strains, and employed a workflow that we named LiQuoR (Liquid Quantification of Rivalry).

Briefly, LiQuoR comprises mixing the attacker and luminescent prey strains in a black 96-well plate with a clear bottom, then incubating them with shaking under the desired conditions for competition. Concomitantly, to evaluate competition in the same attacker-prey strain mixtures on a solid surface, small equal volumes of the bacterial mixtures are taken from the 96-well plate using a floating pin multi-blot replicator and spotted onto a single-well plate containing solid agar media. The single-well plate is also incubated under the desired conditions for competition. Prey viability during competition in liquid media or on solid surface conditions is assessed based on luminescence readings obtained using a microplate reader (Fig. 1D).

We used LiQuoR to determine whether luminescence (relative luminescence units, RLU) can replace CFU counts as a reporter of prey viability during competition. Indeed, RLU showed a similar trend of T6SS-mediated antagonism toward an Awe1-sensitive *Aj* prey (*Aj*Δ*awei1*), on agar and in liquid media, as did CFU counts performed on the same samples used for LiQuoR (Fig. 1E-F; note that the RLU counts were not normalized to account for the smaller volumes used compared to CFU counts). Therefore, we decided to use RLU counts to evaluate prey viability in subsequent competition assays.

We then used LiQuoR to determine whether the ability of *Aj* to use T6SS for kin intoxication in liquid media is restricted to the effector Awe1. By monitoring the viability of *Aj* prey strains sensitive to each of the other three known T6SS effectors Tle1, TseI, and DUF3289 (*Aj*Δ*tlei1, Aj*Δ*tseiI, Aj*Δ*duf3289i*, respectively), we confirmed that all sensitive prey strains are intoxicated by the parental *Aj* strain in a T6SS-dependent manner in liquid media (Supplementary Fig. S2). These results indicate that all *Aj* T6SS effectors are injected into prey cells during competition in liquid media.

### Aeromonas jandaei uses T6SS to kill non-kin prey in liquid media

Following the observations that *Aj* employs T6SS to antagonize kin competitors in liquid media, we asked whether it could do the same against non-kin rivals. Therefore, we employed the LiQuoR method, using *E. coli* as prey during competition against *Aj* in liquid media. As with kin prey, *Aj* antagonized *E. coli* prey in a T6SS-dependent manner (Fig. 2A), indicating that *Aj* can use its T6SS to antagonize both kin and non-kin prey in liquid media.

**Figure 2.**
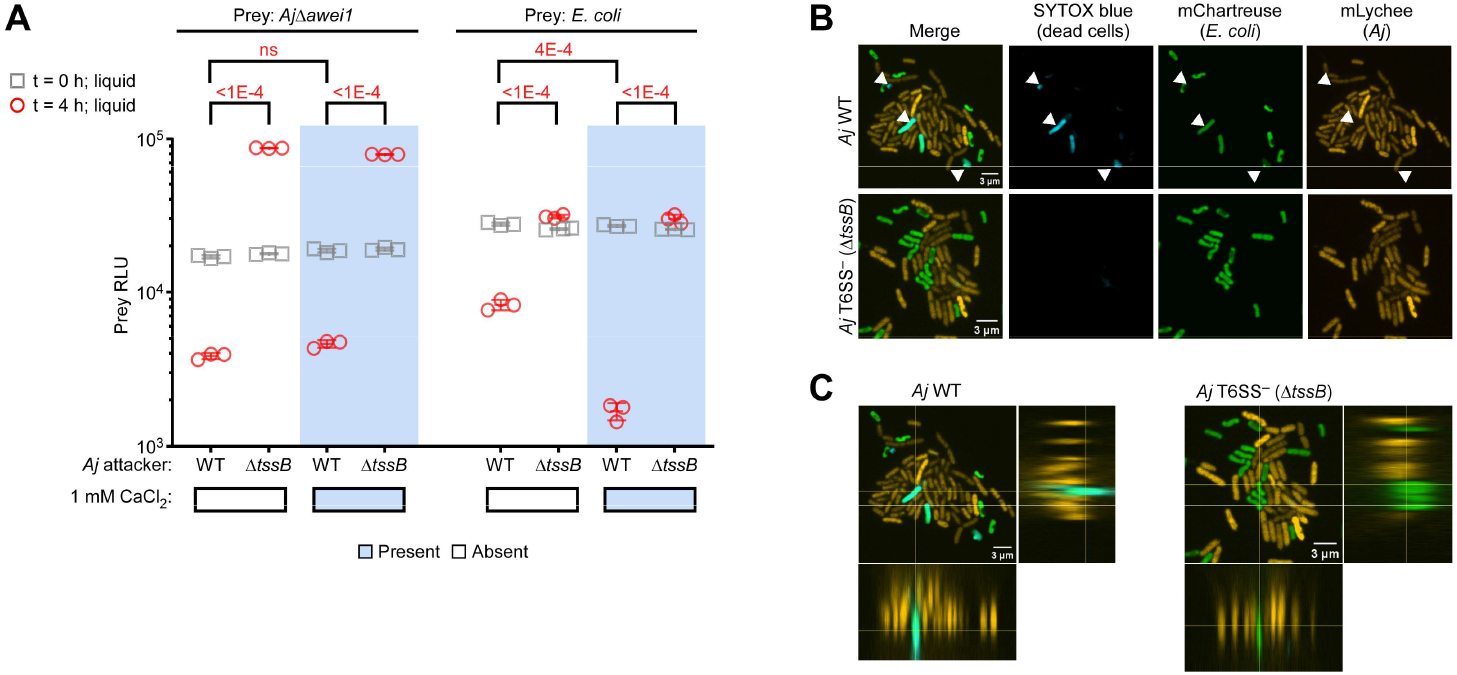
*Aeromonas jandaei* forms aggregates to kill non-kin prey in liquid media. **(A)** LiQuoR-based viability measurements (RLU) of *Aj*Δ*awei1* and *E. coli* prey before (t = 0 h) and after (t = 4 h) co-incubation with the indicated *Aj* attacker at a 4:1 (attacker:prey) ratio in LB with or without 1 mM CaCl_2_ at 30^°^C. Data are shown as the mean ± SD; n = 3 samples. The statistical significance between samples at the 4 h time point for each prey strain was calculated using one-way ANOVA with Tukey multiple comparisons test; ns, P > 0.05. The experiment was repeated three times with similar results. Results from a representative experiment are shown. **(B)** Representative confocal fluorescence microscope images of *Aj* and *E. coli* bacteria co-incubated in LB media containing 1 mM CaCl_2_ at 30^°^C. Triangles denote dead *E. coli* cells. **(C)** Orthogonal view of stacked confocal microscope images shown in B. WT, wild-type. Bar = 3 µm.

Our results imply that *Aj* can form stable cell-cell interactions with prey cells when grown in liquid media. Previous reports have indicated that the presence of calcium in the media can favor bacterial aggregation, thereby increasing cell-cell interactions^53,61^. Since calcium is present in the intestinal tract of fish that are potential *Aeromonas* hosts^62,63^, as well as in brackish waters^64^ from which *Aeromonas* are widely isolated^65,66^, we asked whether adding calcium to the media can enhance the T6SS-mediated antibacterial activity of *Aj*. Interestingly, the addition of CaCl_2_ to the liquid media did not affect the T6SS-mediated killing of kin prey, but it increased the killing of non-kin *E. coli* prey (Fig. 2A). Therefore, we decided to perform subsequent competition assays against *E. coli* prey in liquid media supplemented with CaCl_2_.

Next, we set out to visualize the cell-cell attachment and prey intoxication during co-incubation of *Aj* attackers (constitutively expressing the red fluorescent protein mLychee^67^) and *E. coli* prey (constitutively expressing the green fluorescent protein mChartreuse^67^) in liquid media containing CaCl_2_. As expected, confocal fluorescence microscope imaging revealed mixed attacker-prey aggregates in the bacterial culture after fixation (Fig. 2B-C).

Furthermore, permeabilized *E. coli* prey cells were visible in the presence of wild-type *Aj* attackers, but not in the presence of T6SS^−^ (Δ*tssB*) attackers (as evidenced by the signal of SYTOX blue, a nucleic acid stain that cannot penetrate uncompromised cell membranes) (Fig. 2B-C). Taken together, our results reveal that *Aj* can stably bind to competing bacteria in liquid media and antagonize them using its T6SS.

### Adhesins facilitate T6SS-mediated kin and non-kin killing in liquid media

The observed contact-dependent, T6SS-mediated intoxication of kin and non-kin prey in liquid media led us to hypothesize that *Aj* employs a molecular tool, i.e., adhesin^68^, to attach to prey cells. Such an adhesin could be a large, secreted protein anchored to the cell surface. While examining the *Aj* genome (accession NZ_CP149571.1), we identified three putative adhesins containing repeat sequences and an N-terminal retention_LapA module (Fig. 3A), likely secreted by the type I secretion system (T1SS): (1) WP_225628403.1, which we named CraAj; (2) WP_339058648.1, which we named AdhAj; and (3) WP_339058677.1, which we named LapAj. When examining the latter, we noticed that homologous proteins encoded by other *Aeromonas* genomes are annotated as longer proteins with additional choice-of-anchor K and COG2931 domains at the C-terminus (e.g., WP_440012114.1). Indeed, the gene annotated downstream in *Aj*, predicted to encode WP_339058676.1, contains these two domains, suggesting a possible mistake in the genome sequence that split the adhesin into two genes. Sanger sequencing of a PCR-amplified region connecting the genes encoding WP_339058677.1 and WP_339058676.1 from isolated *Aj* genomic DNA (Supplementary File S1) confirmed a mistake in the genomic sequence found in the RefSeq database (NZ_CP149571.1), resulting in an erroneous early stop codon due to a one-nucleotide insertion. Therefore, the complete amino acid sequence of LapAj comprises the sequences of both WP_339058677.1 and WP_339058676.1 (Fig. 3A and Supplementary File S2).

**Figure 3.**
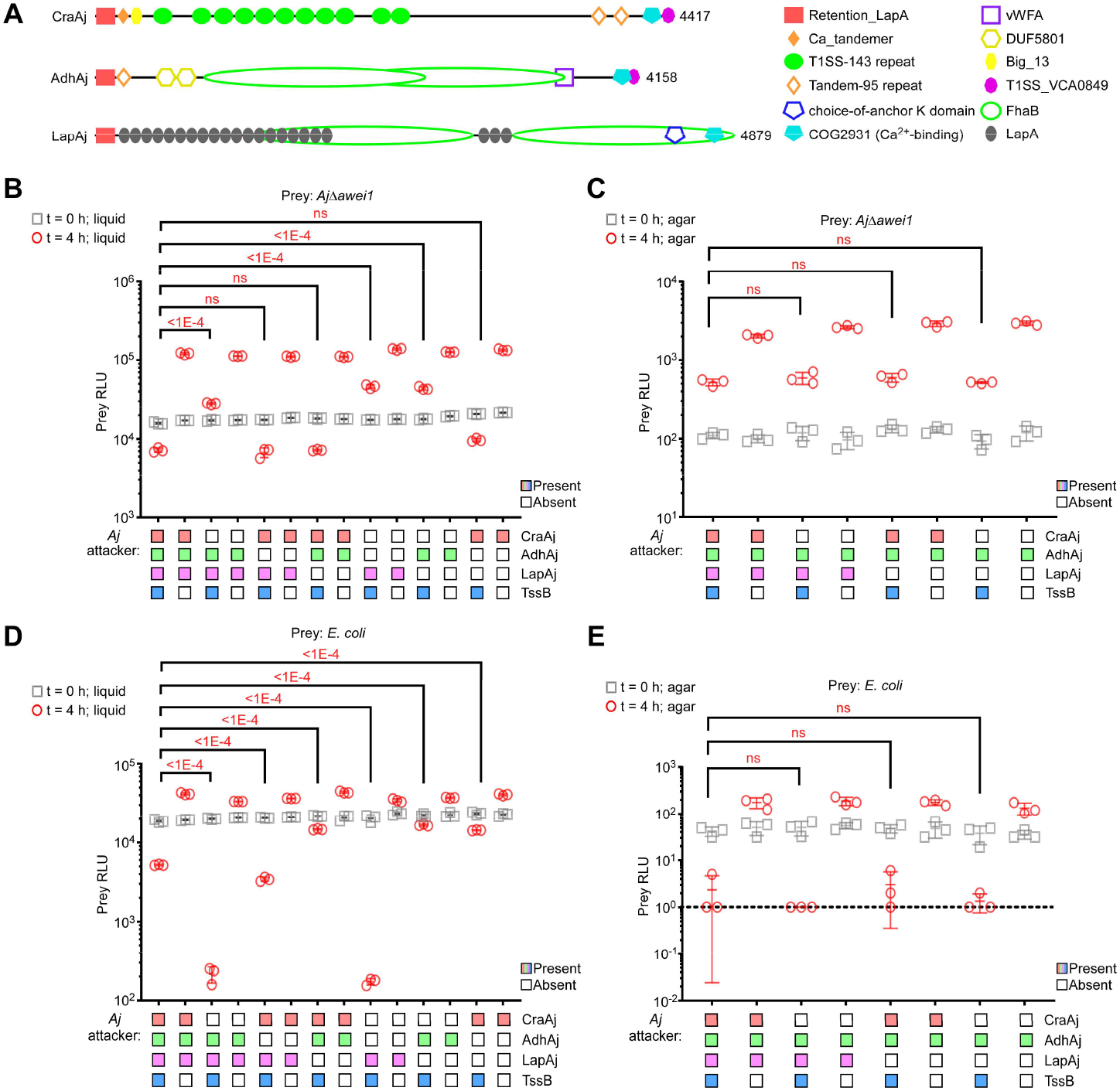
Adhesins have different effects on kin and non-kin T6SS-mediated competition in liquid media. **(A)** Schematic representation of the identified *Aj* adhesins and their domains. **(B-E)** LiQuoR-based viability measurements (RLU) of *Aj*Δ*awei1* (B-C) or *E. coli* (D-E) prey before (t = 0 h) and after (t = 4 h) co-incubation with the indicated *Aj* attacker at a 4:1 (attacker:prey) ratio in LB liquid media with (D) or without (B) 1 mM CaCl_2_ or on LB agar plates (C, E) at 30^°^C. Data are shown as the mean ± SD; n = 3 samples. The statistical significance between mutant and WT samples at the 4 h time point was calculated using one-way ANOVA with Dunnett’s multiple comparisons test; ns, P > 0.05. The experiment was repeated three times with similar results. Results from a representative experiment are shown. WT, wild-type. A dashed line denotes the assay detection limit.

To determine whether CraAj, AdhAj, or LapAj plays a role in the T6SS-mediated competition in liquid media, we constructed strains in which each adhesin was deleted individually, as well as combinations of two deleted adhesins. When these strains were used as attackers in LiQuoR against sensitive kin prey (*Aj*Δ*awei1*), the deletion of *craAj*, either alone or in combination with another adhesin, impaired T6SS-mediated intoxication (Fig. 3B). However, *craAj* deletion did not affect the competition on an agar plate (Fig. 3C), indicating that CraAj plays a role in kin intoxication in liquid media.

Remarkably, when we tested the same *Aj* attacker strains using LiQuoR against non-kin *E. coli* prey, the deletion of *craAj* did not hinder T6SS-mediated intoxication but rather enhanced it (Fig. 3D). Conversly, the deletion of *lapAj*, either alone or in combination with another adhesin (including *craAj*), hindered the T6SS-mediated intoxication of the non-kin prey (Fig. 3D). Notably, neither *craAj* nor *lapAj* deletion affected competition against *E. coli* prey on an agar plate (Fig. 3E). These results suggest that LapAj plays a role in non-kin intoxication in liquid media.

Notably, the deletion of *craAj* or *lapAj* did not affect the growth of *Aj* attackers (Supplementary Fig. S3A-B) or the activity of their T6SS, as determined by monitoring the secretion of the hallmark T6SS component Hcp^58^ (Supplementary Fig. S3C). Furthermore, we did not detect swimming motility in *Aj* (Supplementary Fig. S3D), indicating that adhesin deletions did not affect the ability of *Aj* attackers to swim in liquid media. Taken together, our results demonstrate that in liquid media, the adhesins CraAj and LapAj facilitate T6SS-dependent kin and non-kin intoxication, respectively. Therefore, we hypothesized that CraAj mediates auto-aggregation with kin cells, whereas LapAj mediates co-aggregation with non-kin cells in liquid media.

### CraAj facilitates auto-aggregation

To determine whether CraAj plays a role in *Aj* auto-aggregation, we examined whether its deletion impairs the formation of cell aggregates in liquid media. To this end, we transferred liquid cultures of wild-type, T6SS^−^ (Δ*tssB*), Δ*craAj*, and Δ*lapAj Aj* strains, constitutively expressing the red fluorescent protein mLychee from a plasmid for improved cell detection, into a 96-well plate with LB media. The plate was placed in a microscope (Incucyte) and kept motionless for 5 minutes to allow cell aggregates to settle at the bottom of the well. Then, images were taken (t = 0 h), and the plate was transferred to shake in a 30^°^C incubator for 50 minutes, before images were taken again (t = 1 h) as described above (Fig. 4A). In agreement with our hypothesis, the deletion of *craAj* dramatically reduced the number of cell aggregates detected at the bottom of the well after one-hour of incubation (Fig. 4B-C), indicating that CraAj plays a role in auto-aggregation in liquid media. Notably, the five-minute incubation period without motion before imaging was sufficient for all cell aggregates to settle to the bottom of the well (Supplementary Fig. S4).

**Figure 4.**
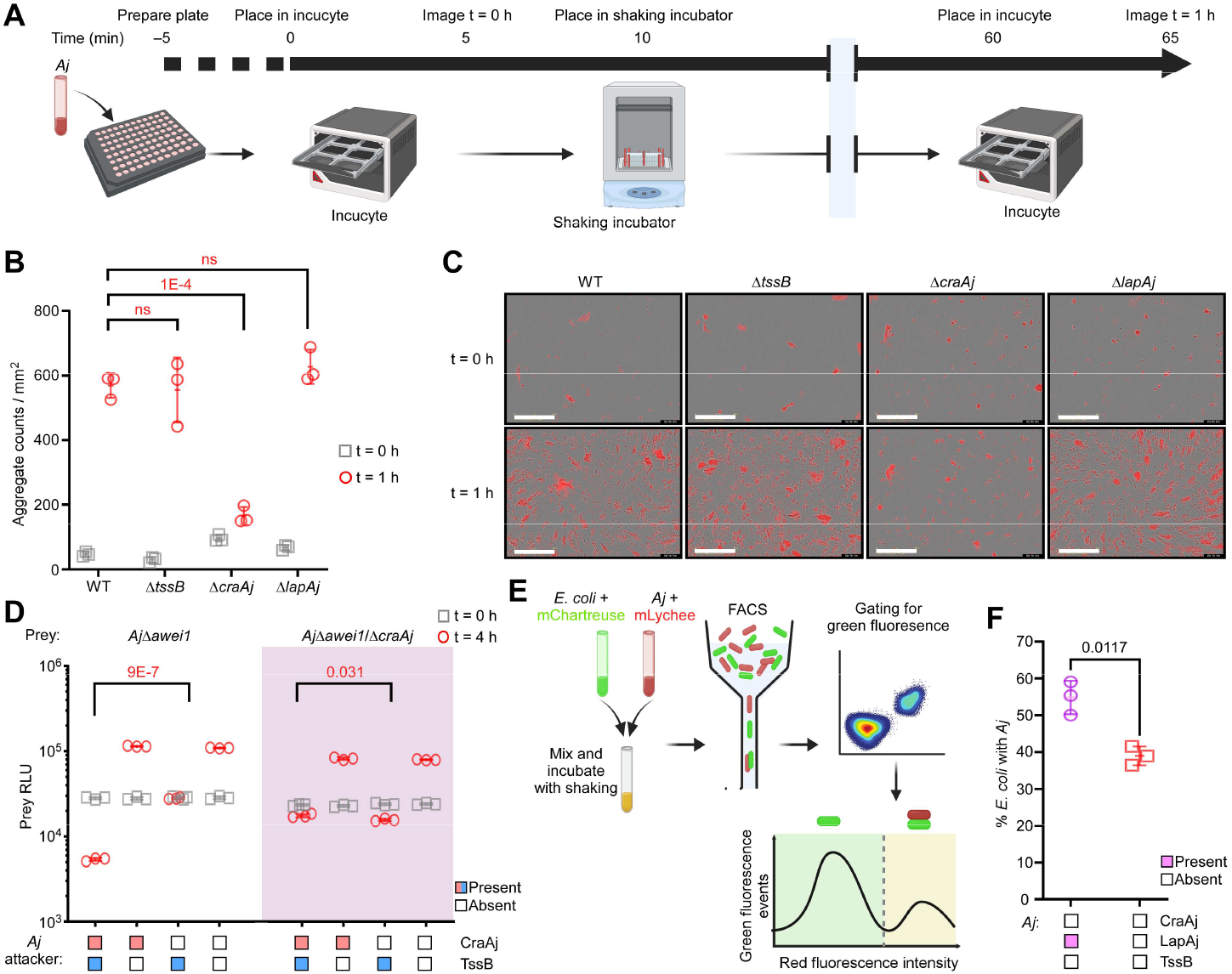
CraAj mediates auto-aggregation and LapAj mediates co-aggregation in liquid media. **(A)** Schematic representation of the experimental workflow used for the data shown in B-C. **(B)** Quantification of *Aj* aggregates from microscope images. The statistical significance between samples at the 1 h time point was calculated using one-way ANOVA with Dunnett’s multiple comparisons test; ns, P > 0.05. **(C)** Representative microscopy images of the indicated *Aj* strains. Merged red fluorescence (mLychee constitutively expressed from a plasmid) and bright field channels are shown. Bar = 400 µm. **(D)** LiQuoR-based viability measurements (RLU) of the indicated *Aj* prey strains before (t = 0 h) and after (t = 4 h) co-incubation with the indicated *Aj* attacker at a 4:1 (attacker:prey) ratio in LB liquid media at 30ºC. The statistical significance between samples at the 4 h time point was calculated using an unpaired, two-tailed Student’s *t* test. **(E)** Schematic representation of the experimental workflow used for the data shown in F. **(F)** The percentage of *E. coli* (mChartreuse; green) signals that are identified in FACS analysis together with *Aj* (mLychee; red) signals. The statistical significance was calculated using an unpaired, two-tailed Student’s *t* test. In B, D, and F, data are shown as the mean ± SD; n = 3 samples. The experiments were repeated three times with similar results. Results from a representative experiment are shown.

We reasoned that the reduced ability to auto-aggregate in the absence of CraAj can explain the observed impairment of kin intoxication by a Δ*craAj* attacker strain (Fig. 3B). Notably, in the kin competition assays described in Fig. 3B, the prey strains maintain their CraAj. We posited that the prey’s CraAj may mediate some level of attachment with the Δ*craAj* attacker, providing a possible explanation for the remaining T6SS-mediated intoxication observed even when CraAj is absent in the attacker strain (Fig. 3B). To investigate this possibility, we repeated the kin intoxication assay with a sensitive prey strain lacking CraAj (*Aj*Δ*awei1*/Δ*craAj*). Although the deletion of *craAj* in the prey strain impaired its intoxication by a wild-type attacker, mirroring the effect of *craAj* deletion in the attacker when competed against a prey containing CraAj, the deletion of *craAj* in both the attacker and prey did not further impair kin intoxication (Fig. 4D). These results imply that CraAj participates in homophilic interactions, in which CraAj from the attacker interacts with CraAj from the prey to facilitate cell-cell interaction in liquid media. Therefore, auto-aggregation and T6SS-mediated intoxication are similarly impaired by the loss of CraAj from either the attacker strain, the prey strain, or both.

### LapAj facilitates co-aggregation with E. coli

To determine whether LapAj plays a role in *Aj* co-aggregation with *E. coli*, we examined whether its deletion impairs cell-cell interactions between the two bacterial strains in liquid media. To this end, we mixed *E. coli* cells expressing the green fluorescent protein mChartreuse with *Aj* cells expressing the red fluorescent protein mLychee. Notably, to prevent *E. coli* intoxication by *Aj* and to enable *Aj* to primarily interact with *E. coli* rather than form auto-aggregates, we used an *Aj* strain lacking CraAj and an active T6SS (Δ*craAj*/Δ*tssB*) as the parental strain. The bacterial mixture was incubated with shaking in LB media containing CaCl_2_, and then analyzed by flow cytometry to identify events in which *E. coli* pass through the interrogation point alone (only green) or attached to *Aj* (green and red) (Fig. 4E and Supplementary Fig. S5). As shown in Fig. 4F, the deletion of *lapAj* in *Aj* significantly reduced the percentage of *E. coli* cells in contact with *Aj* cells, thus confirming a role for LapAj in attachment to *E. coli* cells.

Collectively, our results point to a model in which *Aj* employs at least two adhesins to specifically bind other bacteria in liquid environments. CraAj facilitates auto-aggregation, and LapAj facilitates co-aggregation, thus enabling *Aj* to distinguish kin and non-kin competitors in liquid environments. If a bound cell is sensitive to the effectors injected by its T6SS, *Aj* can intoxicate it (Fig. 5).

**Figure 5.**
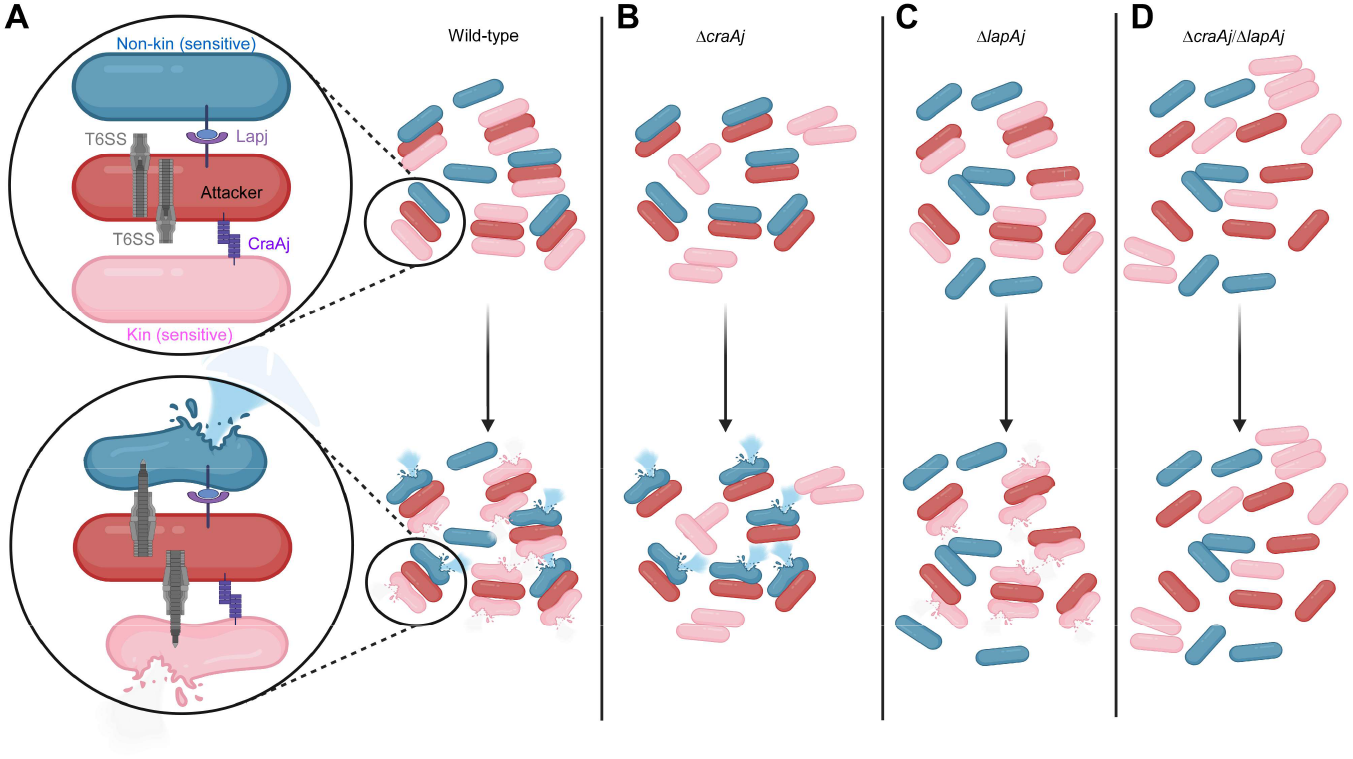
Proposed model for adhesin-mediated intoxication of kin and non-kin competitors in liquid environments. **(A)** wild-type *Aj* contain both CraAj and LapAj, and can bind both kin and non-kin prey. Prey sensitive to *Aj*’s T6SS effectors will be intoxicated. **(B)** In the absence of CraAj, *Aj* is impaired in binding kin cells and becomes more available to bind non-kin cells via LapAj. **(C)** In the absence of LapAj, *Aj* is impaired in binding non-kin cells. **(D)** In the absence of both CraAj and LapAj, *Aj* is impaired in binding both kin and non-kin cells.

### Additional marine bacteria employ T6SS to kill rivals in liquid media

Based on our results and on previous reports of marine bacteria that inflict T6SS-mediated intoxication in liquid media^53–55^, we hypothesized that T6SS-mediated intoxication of rivals in liquid should be a common phenomenon in marine bacteria. In light of possible effects of environmental conditions and cues on T6SS activation and T6SS-mediated competition, we reasoned that different bacteria employ T6SS for competition in liquid under specific conditions; furthermore, it is likely that the identity of the competing prey strain affects the results of the competition. To investigate the possibility of other marine bacteria intoxicating competitors in liquid media using T6SS, we employed two *Vibrio* species whose T6SSs we have previously characterized: (1) the coral pathogen *Vibrio coralliilyticus* ATCC BAA-450 (*Vcor*), which harbors one antibacterial T6SS and one anti-eukaryotic T6SS^12^; and (2) the human pathogen *Vibrio parahaemolyticus* RIMD 2210633 (*Vpara*), which harbors two antibacterial T6SSs^69,70^.

Using the LiQuoR workflow, we competed these *Vibrio* attacker species, either wild-type or inactive in both T6SSs, against *E. coli* prey. The competition was monitored after four hours of co-incubation at 30^°^C in liquid media containing either 1% or 3% (w/v) NaCl, with or without 1 mM CaCl_2_. Remarkably, we found that *Vcor* killed the *E. coli* prey in a T6SS-dependent manner specifically in media containing 3% NaCl; interestingly, the presence of CaCl_2_ in the media slightly impaired the toxic effect against *E. coli* prey, as opposed to its positive effect on the ability of *Aj* to intoxicate *E. coli* in liquid media (Fig. 6A). Conversely, *Vpara* did not intoxicate *E. coli* prey using its T6SSs in liquid media under the tested conditions (Fig. 6B). Taken together, these results indicate that diverse marine bacteria can, under specific conditions, intoxicate rival bacteria in liquid media in a T6SS-dependent manner.

**Figure 6.**
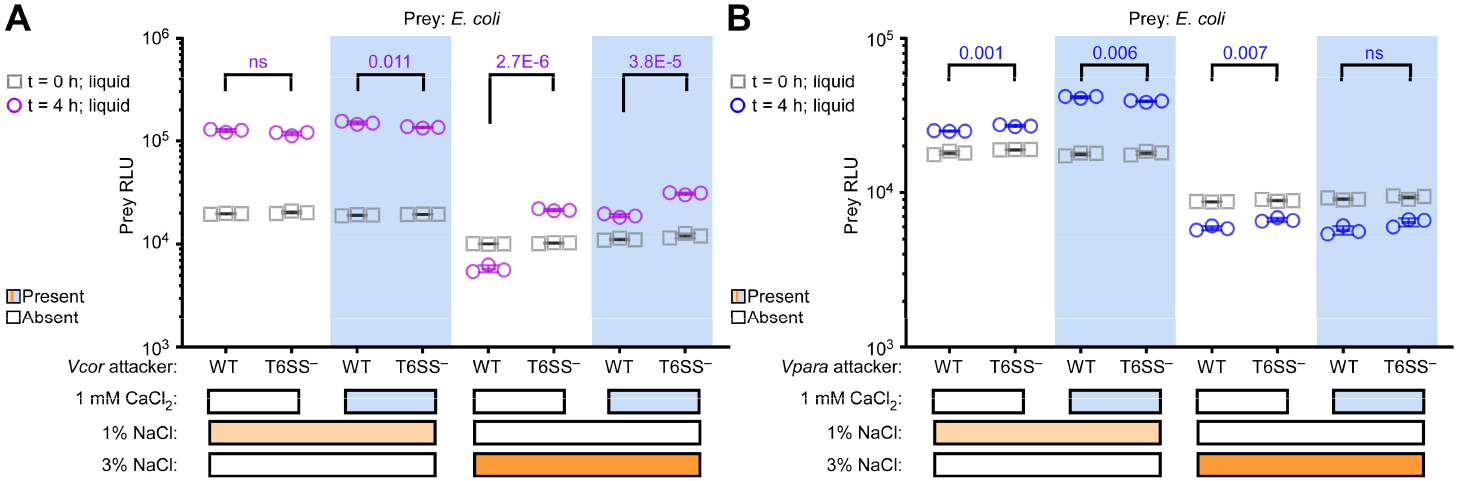
*Vibrio coralliilyticus* use T6SS to intoxicate *E. coli* prey in liquid media. LiQuoR-based viability measurements (RLU) of *E. coli* prey before (t = 0 h) and after (t = 4 h) co-incubation with wild-type (WT) or T6SS^−^ (Δ*hcp1*/Δ*tssM2*) *Vibrio coralliilyticus* ATCC BAA-450 attacker strains **(A)**, or with WT or T6SS^−^ (Δ*hcp1*/Δ*hcp2*) *Vibrio parahaemolyticus* RIMD 2210633 attacker strains **(B)** at a 4:1 (attacker:prey) ratio in LB or MLB media (1% or 3% [w/v] NaCl, respectively) with or without 1 mM CaCl_2_ at 30^°^C. Data are shown as the mean ± SD; n = 3 samples. The statistical significance between samples at the 4 h time point was calculated using an unpaired, two-tailed Student’s *t* test. ns, P > 0.05. The experiments were repeated three times with similar results. Results from a representative experiment are shown.

## DISCUSSION

Since the early days of its discovery, the T6SS has been considered a contact-dependent machine that delivers effectors into neighboring cells only on solid surfaces, where prolonged cell-cell attachment is forced^51^. This notion recently began to change, as three reports suggested that bacteria can engage in T6SS-mediated competition in liquid environments if they can form stable cell-cell attachments under these conditions^53–55^. Here, we show that this phenomenon is likely more common in marine bacteria than previously appreciated, and that bacteria can distinguish kin from non-kin competitors in liquid media.

Bacteria appear to employ diverse mechanisms to mediate cell-cell attachment that facilitates T6SS-dependent antagonism in liquid media. *V. cholerae* was shown to use type IV pili for intra-species interactions^54^, whereas *Aureispira* was proposed to use T9SS for inter-species interactions^55^. Here, we identify a third mechanism employed by *Aj* for both intra- and inter-species interactions. We demonstrate that adhesins play distinct roles in binding kin and non-kin prey, and that the preference for a prey type may be governed by environmental conditions and cues (e.g., the presence or absence of calcium). This suggests a possible ability to direct aggregation (i.e., auto- or co-aggregation) and thus T6SS-mediated aggression toward specific competitors within a niche. However, it is also possible that such changes in aggregation preferences will negatively affect the bacterial population. For example, if loss of auto-aggregation increases non-kin interactions (as we observe when *craAj* is deleted in *Aj*; Fig. 3), the bacterial population could be at risk if the neighboring non-kin competitor has a superior aggression mechanism^71^.

Ting et al. previously engineered a synthetic mechanism enabling T6SS-mediated competition in liquid media^52^. By constructing T6SS-wielding bacteria that present antigen-specific nanobodies on the cell surface (programmed inhibitor cells; PICs), they achieved targeted prey depletion in liquid media. Our results suggest that PICs essentially mimic *Aj*’s natural adhesin-mediated ability to interact with specific prey in liquid media.

Therefore, we propose that natural target-specific adhesins, which facilitate cell-cell interactions in liquid media, are potential tools for identifying prey-specific ligands and for engineering PICs. Future work will reveal the ligand bound by the *Aj* adhesins on kin and non-kin cells and determine the molecular mechanisms that govern *Vcor*-mediated cell-cell interactions in liquid media.

Notably, T6SS-mediated competition was not completely abolished in the absence of CraAj or LapAj, suggesting that *Aj* harbors additional mechanisms that facilitate cell-cell interactions in liquid media. Nevertheless, the requirement for mechanisms mediating prolonged cell-cell attachment reveals a possible non-immunity protein-mediated defense mechanism^48,71,72^ against T6SS antagonism in liquid media. By mutating or losing a receptor or a ligand recognized by the adhesin of a potential attacker, a prey cell can escape prolonged cell-cell contact in liquid media and thus evade T6SS-mediated intoxication (e.g., the partial immunity of *Aj* prey strains lacking CraAj; Fig. 4D).

The species shown to engage in T6SS-mediated competition in liquid media to date, i.e., *Aj, Vcor, V. cholerae, V. fischeri*, and *Aureispira*, are all marine bacteria. This suggests that T6SS-mediated antagonism in liquid media may be an overlooked, widespread phenomenon within this group of bacteria. Importantly, our results imply that one must consider diverse media, growth conditions, and even the bacterial prey species when investigating the potential of T6SS-mediated intoxication in liquid media. In this context, although we did not identify T6SS-mediated intoxication in liquid by *Vpara*, we tested only a limited range of conditions and used only *E. coli* as potential prey. Therefore, it remains possible that *Vpara* employs its T6SS for interbacterial antagonism in liquid under specific conditions.

Finally, we establish LiQuoR as a methodology enabling real-time monitoring of interbacterial intoxication in liquid media and on agar plates simultaneously. LiQuoR eliminates the negative aspects of the canonical CFU counting method on selective antibiotic plates (i.e., labor-intensive, time-consuming, and potentially inaccurate^59^). It also provides a feasible and efficient workflow for screening multiple conditions and prey strains to identify T6SS-mediated intoxication in liquid media in other marine bacteria of interest.

## MATERIALS AND METHODS

### Strains and media

A complete list of bacterial strains used in this study is available in Supplementary Table S1. *Aeromonas jandaei* DSM 7311 (*Aj*) and derivative strains were grown in Lysogeny broth (LB; 1% [w/v] tryptone, 0.5% [w/v] yeast extract, and 1% [w/v] NaCl) or on LB agar plates (1.5% [w/v]) at 30°C. The media were supplemented with kanamycin (30 μg/mL) or chloramphenicol (10 μg/mL) when plasmid maintenance was required. Ampicillin (100 μg/mL) was also used in the media since *Aj* is naturally resistant to it. For induction of gene expression from pBAD-based plasmids, 0.05% (w/v) L-arabinose was added to the media.

*Escherichia coli* strains were grown in 2xYT broth (1.6% [w/v] tryptone, 1% [w/v] yeast extract, and 0.5% [w/v] NaCl) or on LB agar plates at 37°C. The media were supplemented with chloramphenicol (10 μg/mL) or kanamycin (30 μg/mL) to maintain plasmids when required.

*Vibrio parahaemolyticus* RIMD 2210633 (*Vpara*) strains were grown in Marine Lysogeny broth (MLB; LB containing 3% [w/v] NaCl) or Marine Minimal Media (MMM) agar plates (2% [w/v] NaCl, 0.4% [w/v] galactose, 5 mM MgSO_4_, 7 mM K_2_SO_4_, 77 mM K_2_HPO_4_, 35 mM KH_2_PO_4_, 2 mM NH_4_Cl, and 1.5% [w/v] agar) at 30°C.

*Vibrio coralliilyticus* ATCC BAA-450 (*Vcor*) strains were grown in MLB media or on GASW-Tris agar plates (20.8 [g/L] NaCl, 0.56 [g/L] KCl, 4.8 [g/L] MgSO_4_·7H_2_O, 4 [g/L] MgCl_2_·6H_2_O, 0.01 [g/L] K_2_HPO_4_, 0.001 [g/L] FeSO_4_·7H_2_O, 2 [g/L] Instant Ocean sea salts, 6.33 [g/L] Tris base [C_4_H_11_NO_3_], 4 [g/L] tryptone, 2 [g/L] yeast extract, 0.2% [v/v] glycerol, and 1.5% [w/v] agar; pH adjusted to 8.3 with HCl) at 30°C.

### Plasmid and deletion strain construction

A complete list of the plasmids used in this study is available in Supplementary Table S2. For plasmid construction, DNA fragments were amplified by PCR from the relevant genomic DNA or plasmid template, and then assembled using the Gibson assembly method^73^.

To generate chromosomal deletion strains, in-frame deletions were constructed using the *sacB*-based suicide plasmid pDM4^74^. To this end, approximately 1 kb regions upstream and downstream of the target gene(s) were amplified from the genomic DNA of *Aeromonas jandaei* DSM 7311 and cloned into the multiple cloning site (MCS) of pDM4.

All plasmids were first introduced into *Escherichia coli* DH5α (λ-pir) by electroporation and then, when required, transferred into *Aj* via conjugation using *E. coli* HB101 carrying the helper plasmid pRK2013. Transconjugants were selected on agar plates supplemented with appropriate antibiotics; ampicillin was included for *Aj* selection, as it is intrinsically resistant. For allelic exchange using pDM4-based constructs, counter-selection was performed on LB agar plates supplemented with 10% (w/v) sucrose. Successful deletion mutants were verified by PCR. All plasmid constructs were confirmed by sequencing.

### Growth assays

To evaluate the effect of Awe1 expression on *Aj* growth, the indicated *Aj* strains carrying an arabinose-inducible pBAD^K^/*Myc*-His expression plasmid, either empty or encoding Awe1, were grown overnight in LB media supplemented with kanamycin (30 µg/mL). Bacterial cultures were normalized to an OD_600_ of 0.01 in fresh LB media supplemented with kanamycin (30 µg/mL) and transferred in quadruplicate (200 µL per well) into 96-well plates. The bacterial cultures were incubated at 30°C in a BioTek SYNERGY H1 microplate reader with continuous shaking (205 cpm). After 2 h, L-arabinose was added to a final concentration of 0.05% (w/v) to induce expression from the plasmid. OD_600_ measurements were recorded every 10 minutes, and data were analyzed using GraphPad Prism.

To evaluate the effect of chromosomal deletions on bacterial growth, *Aj* strains carrying a plasmid for constitutive expression of the fluorescent protein mLychee were grown overnight and then normalized to an OD_600_ of 0.01 in fresh LB media supplemented with chloramphenicol (10 µg/mL) to maintain the plasmid. Bacterial cultures were transferred in triplicate to a black 96-well plate with a clear bottom (200 µL per well) and incubated at 30°C in a BioTek SYNERGY H1 microplate reader with continuous shaking (205 cpm). OD_600_ and fluorescence (excitation at 568 nm; emission at 596 nm; fluorescence detected from the bottom of the plate) measurements were recorded every 10 minutes, and the data were analyzed using GraphPad Prism.

To evaluate whether T6SS-mediated toxicity in liquid media is contact-independent, *Aj* strains containing either an active (wild-type) or an inactive (ΔtssB) T6SS were grown overnight in LB. Bacterial cultures were then normalized to an OD_600_ of 0.5 in fresh media and incubated for four additional hours at 30°C with shaking. The spent culture supernatants were then centrifuged and filtered through 0.22 µm membranes to remove residual bacterial cells. Cleared supernatants were mixed 1:1 with fresh LB media and used as growth media for Awe1-sensitive *Aj* bacteria (Δ*awei1*), which were inoculated at an initial OD_600_ of 0.01 in triplicate into a 96-well plate (200 µL per well). The plate was incubated at 30°C in a BioTek SYNERGY H1 microplate reader with continuous shaking (205 cpm). OD_600_ measurements were recorded every 10 minutes, and the data were analyzed using GraphPad Prism.

### Protein expression in *Aj*

To monitor the expression Awe1 in different strains, *Aj* strains carrying an arabinose-inducible pBAD^K^/*Myc*-His expression plasmid, either empty or encoding Awe1 with a C-terminal Myc tag, were grown overnight in LB media supplemented with kanamycin (30 µg/mL). Bacterial cultures were normalized to OD_600_ = 0.5 in 3 mL of LB media containing kanamycin (30 µg/mL). Then, the bacterial cultures were incubated at 30°C with shaking (220 rpm) for 1.5 h, followed by induction with 0.05% (w/v) L-arabinose. After an additional 1.5 h of incubation, 0.5 OD_600_ units of cells were harvested. Cell pellets were resuspended in 2x Tris-Glycine SDS sample buffer (Novex, Life Sciences) supplemented with 5% (v/v) β-mercaptoethanol. The samples were incubated at 95°C for 10 minutes and loaded onto Mini-PROTEAN TGX stain-free precast gels (Bio-Rad) for SDS-PAGE. Proteins were transferred onto 0.2 µm nitrocellulose membranes using the Trans-Blot Turbo system (Bio-Rad). Membranes were probed with mouse monoclonal anti-c-*Myc* antibodies (clone 9E10; Santa Cruz Biotechnology, sc-40) at a 1:1,000 dilution. Protein signals were visualized using enhanced chemiluminescence (ECL) and imaged on a Fusion FX6 system (Vilber Lourmat).

### Competition assays

#### CFU-based competition assays

To evaluate interbacterial competition using CFU counts, attacker and prey strains, the latter containing a chloramphenicol-resistance plasmid for selection, were grown overnight in appropriate media. Bacterial cultures were normalized to an OD_600_ of 0.5 and mixed at a 4:1 (attacker:prey) ratio. For liquid media-based competition assays, bacterial mixtures were transferred in triplicate either to glass tubes (3 mL per tube) or to 96-well plates (200 µL per well), as indicated. Glass tubes and 96-well plates were incubated at 30°C with shaking at 220 rpm. Plates were wrapped with Parafilm during incubation. For agar-based competition assays, bacterial mixtures were spotted in triplicate onto LB agar plates (25 µL per spot) and incubated at 30°C.

To evaluate prey viability at the beginning of the experiment (t = 0 h), 25 µL aliquots were collected from all competition mixtures immediately after mixing and added to 975 µL of LB media. Then, the samples were serially diluted (10-fold) and plated on selective LB agar plates supplemented with chloramphenicol (10 µg/mL). Prey CFUs were counted after an overnight incubation at 30°C.

To evaluate prey viability at the end of the experiment (t = 4 h): For liquid media competition assays, 25 µL aliquots were collected from each culture and added to 975 µL of LB medium, serially diluted (10-fold), and plated as described above. For agar-based competition assays, each competition spot was scraped into 1 mL of LB media, serially diluted (10-fold), and plated as described above. Plates were incubated overnight at 30°C, and colonies were counted the following day.

#### RLU-based competition assays (LiQuoR)

To evaluate interbacterial competition using RFU counts, attacker and prey strains, the latter containing the pCmLux2 plasmid for constitutive expression of the *Lux* operon, were grown overnight in appropriate media. Bacterial cultures were normalized to an OD_600_ of 0.5 and mixed at a 4:1 (attacker:prey) ratio. Bacterial mixtures were transferred in triplicate to black, flat, clear-bottom 96-well plates (Greiner, cat. 655090; 200 µL per well). Where indicated, 1 mM CaCl_2_ was added to the media. For agar-based RLU competition assays, bacterial mixtures were sampled from the liquid media-containing 96-well plate using a floating pin multi-blot replicator and spotted onto a single-well plate (Tray plate, SPL Life, cat. 31001) containing LB agar. All competition plates were wrapped in Parafilm and incubated at 30°C for 4 h. Liquid media competition plates were incubated with shaking. Relative luminescence units (RLUs) were measured at the indicated time points using a BioTek SYNERGY H1 microplate reader. Prior to the initial luminescence measurement, the plates were incubated in the microplate reader for 10 minutes at 30°C with shaking to equilibrate the media temperature. Luminescence was recorded in top-view luminescence detection mode, without a plate lid. Negative RLU values obtained after media-only background subtraction were set to 1 for subsequent calculations.

### Confocal fluorescence microscopy

Bacteria were grown overnight in LB media containing 10 µg/mL chloramphenicol at 200 rpm and at 37^°^C for *E. coli* and 30^°^C for *Aj*. In the morning, the cultures were diluted to OD_600_ of 0.5, mixed in a 1:1 ratio and cultured for 3.5 h at 30^°^C and 200 rpm in LB media containing 10 µg/mL chloramphenicol and 1 mM CaCl_2_. The cultures were then fixed with 2% (v/v) PFA for 10 minutes at room temperature, washed once with 1xPBS and imaged on an LB-PBS agarose pad^75^. Bacterial aggregates were imaged using a Nikon ECLIPSE Ti2 microscope with a CrestOptics X-Light V3 spinning disk confocal system, equipped with a Kinetix sCMOS camera (Teledyne) and a 100x oil objective (numerical aperture 1.45) at room temperature, using a Z-step size of 0.2 µm. To visualize permeabilized cells, 0.5 µM SYTOX Blue Nucleic Acid Stain (Invitrogen, S11348) was added to the agarose pad. Images were acquired using the NIS-Elements Version 6 and analyzed using Fiji.

### Auto-aggregation assays

A 96-well plate was prepared with 200 µL of *Aj* strains harboring a plasmid for the constitutive expression of the red fluorescent protein mLychee at OD_600_ = 1 in triplicate. The plate was immediately placed in an Incucyte SX5 system, maintained for 5 minutes (to allow aggregates to sink to the bottom), and images were acquired (t = 0 h). The plate was then transferred to a shaking incubator at 30^°^C for an additional 50 minutes. After incubation, the plate was placed in the Incucyte, maintained for 5 minutes, and images were acquired (t = 1 h). To confirm that 5 minutes is sufficient for all cell aggregates to sink to the bottom of the wells, we also performed this assay when pictures at the 1 h time point were acquired immediately after placing the plate in the Incucyte, as well as 4, 6, 8, and 17 minutes after that time. Image analysis was performed using Incucyte SX5 analysis software. Aggregates were quantified from imaging data, with an aggregate defined as an object in the orange channel (mLychee) with an Orange Count Unit (OCU; i.e., fluorescence intensity) ≥ 50 and an area ≥ 150 µm^2^. The resulting data were exported to GraphPad Prism for further analysis. The assay was performed three times with similar results; representative results are shown.

### Co-aggregation assays

*Aj* strains (Δ*tssB*/Δ*craAj* or Δ*tssB*/Δ*craAj*/Δ*lapAj*), harboring a plasmid for the constitutive expression of the red fluorescent protein mLychee, were grown overnight in LB media with chloramphenicol (10 µg/mL); *E. coli*, harboring a plasmid for the constitutive expression of the green fluorescent protein mChartreuse, were grown overnight in 2xYT media with chloramphenicol (10 µg/mL). Bacterial cultures were normalized to OD_600_ = 0.5 and then mixed at a 4:1 ratio (*Aj*:*E. coli*). Triplicates of these mixtures were transferred into a 96-well plate (200 µL per well) and incubated with shaking at 30°C for 2 h. Following incubation, 20 µL from each well were transferred into tubes containing 1 mL of LB media. Samples were analyzed by ThermoFisher Scientific Attune NxT. A minimum of 100,000 events were acquired per sample and initially gated based on forward scatter (FSC) and side scatter (SSC). mChartreuse fluorescence (*E. coli*) was excited with the 488 nm blue laser and detected using a 530/30 nm filter, while mLychee fluorescence (*Aj*) was excited with the 561 nm yellow laser and detected using a 620/15 nm filter. The mChartreuse-positive (*E. coli*) population was gated for downstream analysis, within which the mLychee-positive (*Aj*) population was quantified (Supplementary Fig. S5). Data were analyzed using FlowJo software and subsequently exported to GraphPad Prism. Co-aggregation was quantified as the percentage of red-positive events within the green-positive population. The assay was performed three times with similar results, and the results from a representative experiment are shown.

### Protein secretion assays

Hcp secretion assays were performed as previously described^58^, with minor modifications. *Aj* strains were grown overnight at 30°C in LB media. Bacterial cultures were normalized to an OD_600_ of 0.5 in 5 mL fresh LB media (supplemented with 1 mM CaCl_2_, where indicated) and incubated for three additional hours at 30°C with shaking (220 rpm). For expression (cell) fractions, cells corresponding to 0.5 OD_600_ units were harvested by centrifugation and resuspended in 2x Tris-Glycine SDS sample buffer (Novex, Life Sciences) supplemented with 5% (v/v) β-mercaptoethanol. For secretion (media) fractions, culture volumes corresponding to 10 OD_600_ units were filtered through 0.22 µm membranes, and proteins were precipitated. Precipitates were washed twice with cold acetone and resuspended in 20 µL of 10 mM Tris-HCl (pH 8.0), followed by the addition of 20 µL of 2x Tris-Glycine SDS sample buffer supplemented with 5% (v/v) β-mercaptoethanol and 0.5 µL of 1 M NaOH. Expression and secretion samples were boiled at 95°C for 10 min and 5 min, respectively, separated on Criterion TGX stain-free precast gels (Bio-Rad), and transferred onto 0.2 µm nitrocellulose membranes using the Trans-Blot Turbo system (Bio-Rad). Membranes were probed with a custom-made anti-Hcp primary antibody (1:1,000; GenScript; polyclonal rabbit antibodies raised against the peptide DPQSGQPAGQRVHKC). Anti-RpoB antibodies (1:40,000; BioLegend, clone 8RB13) were used as a loading and lysis control. Signals were detected by ECL and visualized using the Fusion FX6 system.

### Swimming assays

To evaluate swimming motility, *Aj* strains were grown on LB agar plates, and single colonies were picked and inoculated into LB solidified with 0.3% (w/v) agar (supplemented or not with 1 mM CaCl_2_) by stabbing the agar with a sterile toothpick. Swimming plates were incubated at 30°C for 16 h.

## Supporting information

Supplementary Information (Figs, Tables, and References)

## ACKNOWLEDGEMENTS

We thank members of the Salomon lab for helpful discussions. The project received funding from the Israel Science Foundation (ISF grant number 1362/21 to DS and 2174/22 to MG), and was supported by the National Center of Competence in Research AntiResist funded by the Swiss National Science Foundation (51NF40_180541) to MB. CMF was partially supported by a scholarship from the Clore Israel Foundation. The funders played no role in the study design, data collection and analysis, decision to publish, or preparation of the manuscript. Figures 1D, 4A, 4E, and 5 were prepared using biorender.com.

