## Supplementary File S1 for "*Aeromonas* adhesins facilitate kin and non-kin attachment to enable T6SS-mediated antagonism in liquid"

- The position of the erroneous nucleotide insertion in the NZ_CP149571.1 RefSeq sequence if found between the two bolded nucleotides.

> Forward sequencing

NNNNNNNNNCCCNTGNNCNNCTTTTTATCNCNNCTCTCTACTGTTTCTCCATACCCGTTTTTTTGGGCTAGCGAATTCGAGCTCGGTACCCGGGGATCCTCTAGAAATAATTTTGTTTAACTTTAAGAAGGAGATATACATGTAGTGCTGAGACCGTCATTGTGGACAACGACGCTCCCCCTACCGTGGTGATCAATAAACCTGCGGTGGTGTCCGAAGAGGGGTTGGCCAACGGGTTGCCTGATAACGCAGGTAATACAGATACCACCAATGCGACCGTCGCCAGCGGCACTATTACCGTGGGGGATGTCGATAGCAGTAGCGTGACGGTCAGCCTCTCTG**GT**CCATCCGGACTCAAGTCCGGCGGGCTGGATGTGAAGTGGAACTGGGATGCATCAGGTAAGCTGCTGCTGGTGGTGGTGATTACAAGGATGACGACGATAAGTGAAAGCTTGGCTGTTTTGGCGGATGAGAGAAGATTTTCAGCCTGATACAGATTAAATCAGAACGCAGAAGCGGTCTGATAAAACAGAATTTGCCTGGCGGCAGTAGCGCGGTGGTCCCACCTGACCCCATGCCGAACTCAGAAGTGAAACGCCGTAGCGCCGATGGTAGTGTGGGGTCTCCCCATGCGANAGTAGGGAACTGCCAGGCATCAAATAAAACGAAAGGCTCAGTCGAAAGACTGGGCCTTTCGTTTTTATCTGTTGTTTGTCGGTGAACGCTCTCCTGAGTAGGACAAATCCGCCGGGAGCGGATTTGAAACGTTGCGAAGCAACGGCCCGGAGGGTGGCGGGCAGGACGCCCCGCCATAAACTGCCAGGCATTCAAATTAAGCAGAAGGNCATC

> Reverse sequencing

NNNNNNNNNNCCNAANCNNNNCNNNCTTTCACTTATCGTCGTCATCCTTGTAATCACCACCACCAGCAGCAGCTTACCTGATGCATCCCAGTTCCACTTCACATCCAGCCCGCCGGACTTGAGTCCGGATGG**AC**CAGAGAGGCTGACCGTCACGCTACTGCTATCGACATCCCCCACGGTAATAGTGCCGCTGGCGACGGTCGCATTGGTGGTATCTGTATTACCTGCGTTATCAGGCAACCCGTTGGCCAACCCCTCTTCGGACACCACCGCAGGTTTATTGATCACCACGGTAGGGGGAGCGTCGTTGTCCACAATGACGGTCTCAGCACTACATGTATATCTCCTTCTTAAAGTTAAACAAAATTATTTCTAGAGGATCCCCGGGTACCGAGCTCGAATTCGCTAGCCCAAAAAAACGGGTATGGAGAAACAGTAGAGAGTTGCGATAAAAAGCGTCAGGTAGGATCCGCTAATCTTATGGATAAAAATGCTATGGCATAGCAAAGTGTGACGCCGTGCAAATAATCAATGTGGACTTTTCTGCCGTGATTATAGACACTTTTGTTACGCGTTTTTGTCATGGCTTTGGTCCCGCTTTGTTACAGAATGCTTTTAATAAGCGGGGTTACCGGTTTGGTTAGCGAGAAGAGCCAGTAAAAGACGCAGTGACGGCAATGTCTGATGCAATATGGACAATTGGTTTCTTCTCTGAATGGCGGGAGTATGAAAAGTATGGCTGAAGCGCAAAATGATCCCCTGCTGCCGGGATACTCGTTTAATGCCCATCTGGTGGCGGGTTTAACGCCGATTGAGGCCAACGGTTATCTCGATTTTTTTATCGACCGACCGCTGGGAATGAAAGGTTATATTCTCAATCTCACCATTCGCGGTCANGGGGTGGTGAAAAATCAGGGAACGAGAATTTGNTTGCCGACCGGGTGATATTTTGCTGTTCCCGCCAGGAGAGANTCATCACTACGGTCGTCATCCGGAGGNTCGCGAATGGTATCANCAGTGGGTTTACTTTCGTCCGCGCGNCTACNGGNATGAATGCCTAACTGGNNGTCAANNATTTGCCAATACGGNNCTTTNNNCCCNGATNANCGCCACCANCNGCATTTCANNNACCTGNTTGGNAANTCANNNNNNNNNNNNNNNNNNNNTNNTNNGNNNTNCTGNNNNAATNTGNTTNNNNNNNNNNNNGNNGNNCANGNNNNCNNNTNANCGANNTNNNNNNCNNNNCCANCNNNNNNNNNN
