## Supplementary File S2 for "*Aeromonas* adhesins facilitate kin and non-kin attachment to enable T6SS-mediated antagonism in liquid"

**Corrected LapAj amino acid sequence:**

MITNAITLEQGVTVTQLKGKIYLLAADGSRKLLVEGDVLPKGAVIVSPDGGSFMGGNQSFNVQPAGQPSESAEEGDAPQLAQNGAPGTPSDINALQQAILGGADPTKTLEAPAAGGAPAAGGGGGVGGVAGASGNGGFVTIDRVGDATISQAGFDTTYNNGTAPTLDATGIADQPFISANLTLSAQPQVNEGGSITFTASVDVPVFGTDLVITLSNGAVITIPAGQTTGSVTVAVPADSPYVDPSAITVSITGTQGGGFNQIITGPSTTTQINDTIDTTTVTLGAPAQVNEGGQITYSASVNNAPQSDLVLTLSNGASITIKAGELSGSVTVDAPSDDVYKDGSSLTVSITGSSGGNYEQLDTSSKVTTEVADTVDTTTVTLSAPAQVNEGGQITYSASVNNAPQSDLVLTLSNGASITIKAGELSGSVTVDAPSDDVYKDGSSLTVSITGSSGGNYEQLDTSSKVTTEVADTVDTTTVTLSAPAQVNEGGQITYSASVNNAPQSDLVLTLSNGASITIKAGELSGSVTVDAPSDDVYKDGSSLTVSITGSSGGNYEQLDTSSKVTTEVADTVDTTTVTLSAPAQVNEGGQITYSASVNNAPQSDLVLTLSNGASITIKAGELSGSVTVDAPSDDVYKDGSSLTVSITGSSGGNYEQLDTSSKVTTEVADTVDTTTVTLSAPAQVNEGGQITYSASVNNAPQSDLVLTLSNGASITIKAGELSGSVTVDAPSDDVYKDGSSLTVSITGSSGGNYEQLDTSSKVTTEVADTVDTTTVTLSAPAQVNEGGQITYSASVNNAPQSDLVLTLSNGASITIKAGELSGSVTVDAPSDDVYKDGSSLTVSITGSSGGNYEQLDTSSKVTTEVADTVDTTTVTLSAPAQVNEGGQITYSASVNNAPQSDLVLTLSNGASITIKAGELSGSVTVDAPSDDVYKDGSSLTVSITGSSGGNYEQLDTSSKVTTEVADTVDTTTVTLSAPAQVNEGGQITYSASVNNAPQSDLVLTLSNGASITIKAGELSGSVTVDAPSDDVYKDGSSLTVSITGSSGGNYEQLDTSSKVTTEVADTVDTTTVTLSAPAQVNEGGQITYSASVNNAPQSDLVLTLSNGASITIKAGELSGSVTVDAPSDDVYKDGSSLTVSITGSSGGNYEQLDTSSKVTTEVADTVDTTTVTLSAPAQVNEGGQITYSASVNNAPQSDLVLTLSNGASITIKAGELSGSVTVDAPSDDVYKDGSSLTVSITGSSGGNYEQLDTSSKVTTEVADTVDTTTVTLSAPAQVNEGGQITYSASVNNAPQSDLVLTLSNGASITIKAGELSGSVTVDAPSDDVYKDGSSLTVSITGSSGGNYEQLDTSSKVTTEVADTVDTTTVTLSAPAQVNEGGQITYSASVNNAPQSDLVLTLSNGASITIKAGELSGSVTVDAPSDDVYKDGSSLTVSITGSSGGNYEQLDTSSKVTTEVADTVDTTTVTLSAPAQVNEGGQITYSASVNNAPQSDLVLTLSNGASITIKAGELSGSVTVDAPSDDVYKDGSSLTVSITGSSGGNYEQLDTSSKVTTEVADTVDTTTVTLSAPAQVNEGGQITYSASVNNAPQSDLVLTLSNGASITIKAGELSGSVTVDAPSDDVYKDGSSLTVSITGSSGGNYEQLDTSSKVTTEVADTVDTTTVTLSAPAQVNEGGQITYSASVNNAPQSDLVLTLSNGASITIKAGELSGSVTVDAPSDDVYKDGSSLTVSITGSSGGNYEQLDTSSKVTTEVADTVDTTTVTLSAPAQVNEGGQITYSASVNNAPQSDLVLTLSNGASITIKAGELSGSVTVDAPSDDVYKDGSSLTVSITGSSGGNYEQLDTSSKVTTEVADTVDTTTVTLSAPAQVNEGGQITYSASVNNAPQSDLVLTLSNGASITIKAGELSGSVTVDAPSDDVYKDGSSLTVSITGSSGGNYEQLDTSSKVTTEVADTVDTTTVTLSSATNGQTVTEGGSIVYTASVSNPVTGSPLTVTLSNGVVITIPVGSSSANSDPVPVRGDDIYQQGEELLSVTIDKTSGGNYENVVSTGTITNTVVDDQDLPTLTLTGDANVVEGGKASYTLSISDAPKTDLTVKVVVGHITTDEGDVQAVTRDVVIKAGTTSVSFDVATKDDVYAEPAEQFKVSVSGTSGGGYEKAPALPGAVTTTITDEDGSPDQPKDAPTLTLTGDANVVEGGKASYTLSISDAPKTDLTVKVVVGHITTDEGDVQAVTRDVVIKAGTTSVSFDVATKDDVYAEPAEQFRVSVSGTSGGGYEKAPALPGAVTTTITDEDGSPDQPKDAPTLTLTGDANVVEGGKASYTLSISDAPKTDLTVKVVVGHITTDEGDVQAVTRDVVIKAGTTSVSFDVATKDDVYAEPAEQFRVSVSGTSGGGYEKAPALPGAVTTTITDEDGSPDQPKDAPTLTLTGDANVVEGGKASYTLSISDAPKTDLTVKVVVGHITTDEGDVQAVTRDVVIKAGTTSVSFDVATKDDVYAEPAEQFRVSVSGTSGGGYEKAPALPGAVTTTITDEDGSPDQPKDAPTLTLTGDANVVEGGKASYTLSISDAPKTDLTVKVVVGHITTDEGDVQAVTRDVVIKAGTTSVSFDVATKDDVYAEPAEQFKVSVSGTSGGGYEKAPALPGAVTTTITDEDGSPDQPKDAPTLTLTGDANVVEGGKASYTLSISDAPKTDLTVKVVVGHITTDEGDVQAVTRDVVIKAGTTSVSFDVATKDDVYAEPAEQFRVSVSGTSGGGYEKAPALPGAVTTTITDEDGSPDQPKDAPTLTLTGDANVVEGGKASYTLSISDAPKTDLTVKVVVGHITTDEGDVQAVTRDVVIKAGTTSVSFDVATKDDVYAEPAEQFRVSVSGTSGGGYEKAPALPGAVTTTITDEDGSPDQPKDAPTLTLTGDANVVEGGKASYTLSISDAPKTDLTVKVVVGHITTDEGDVQAVTRDVVIKAGTTSVSFDVATKDDVYAEPAEQFRVSVSGTSGGGYEKAPALPGAVTTTITDEDGSPDQPKDAPTLTLTGDANVVEGGKASYTLSISDAPKTDLTVKVVVGHITTDEGDVQAVTRDVVIKAGTTSVSFDVATKDDVYAEPAEQFKVSVSGTSGGGYEKAPALPGAVTTTITDEDGSPDQPKDAPTLTLTGDANVVEGGKASYTLSISDAPKTDLTVKVVVGHITTDEGDVQAVTRDVVIKAGTTSVSFDVATKDDVYAEPAEQFRVSVSGTSGGGYEKAPALPGAVTTTITDEDGSPDQPKDAPTLTLTGDANVVEGGKASYTLSISDAPKTDLTVKVVVGHITTDEGDVQAVTRDVVIKAGTTSVSFDVATKDDVYAEPAEQFKVSVSGTSGGGYEKAPALPGAVTTTITDEAVPGAEDTVYAQIEVDQSSVVEGGALKYTVSLVDKDGNAVTLPAGKVVTLNLAWSGAAANDGDTSGRPASVTIGANGKAEFTVTTVNDTIYEMDEKLAVTISGVKINSAFEAISIVNGKGSAETVIVDNDAPPTVVINKPAVVSEEGLANGLPDNAGNTDTTNATVASGTITVGDVDSSSVTVSLSGRSGLKSGGLDVKWNWDASGKVLTGYTGTLGGADYKAVLDVKLTAPAGSTKGDWGYDVTLKGALDHPVTGAEDTLSFNVGVSVSDGQSTTVGSLPITVEDDAPIAGDMAAVSLVKTSIPDVLTGKFSLTGYVGDSGSIDAGKFTITARGFESSTSSRLVDAKVNASVDGLGVKSTSAPYHNIDNEIDFRKFADGSSVSEELVIKLDAGTVAYGAKIEFSKIYGAELECGVVEFWRDGKLVATQTFSSNAANGDYAANFQVQQGGFDTMVIKATDNGKSASWGDNSDFTVKSIEFLGSSTPQAIAYGSGTLDPQWGADGKGRFELVTGTVESGLKTVAGLAIAITADGANTLLGKDSNGSLIFKMEFTPATGKWEFFQYAEMQRPADGDIDFTFKAYDRDGDGSQGSFAVNPLVRPDVMGVSNASVTEGGALQHVVTLSGATNVATEYSLSIAGSGAHAASSADWGALQFTNGVTYNSTTGKITVPAGVSSFTVSLNTVNDTLPEFTETLAISVGGVSGTGTIVDNDLSVRLGVGLVDEDGLSGGNAGLPEPVGAPTSGPLSVTQSLQVTDGNGSNVSGVQLKLVSITGLSGITGIDGQPVNVVQDGAGLKGYFGTNPANVAFTVTVDNNSNPPSYTFTLIKPLSHLVDGQSSVLSSQDELRLTVNYEVSKAGSDTASGSFDVAVRDDVPVAVVDDATLNVVVDSFQFSGVEASWINVVGGKNLEYFDGPDNDSANDQIRWGGKNEVKSGYGFADNDAALNGQIPLNEEIKLGTFTHYNYPISSGTSISAVTMKAIFSVTDSMGRVTPVTVTINFNHNETPNDGADPRDIITIGTATASFNFEGKVYSLEVLGFRDTGGNVVKTIYTNENASNSFDLIVKLVEGSGYQLPQTTGNVLANDLAGADGGLSVIGYGVGSSSTTYTNGAGTQVVGLYGVLTILANGAYTYQVTKNGSQIPADAREVFSYSVRDGDGDTTNSTLTISVNPVDSNGVPVHLPLTVDGTDLNDSIVVRNGENAANPDRLDVAFGGNLLGEVTTSNGGTDHVHTGISYNKGSLDQVVSGGAGNDHIETGSGNDVIYAGKTGADGFGTDDSLQLTVAQLKAHHIMTGSLSGTDAMLDGDGLLLSVDVSSSKADVVNGGSGNDKIYGQSGSDILFGGTGDDYIDGGSHNDALRGGLGNDILIGGLGNDVMRGDGGSDTFVWNQGDTVSGSLTKDYIMDFNKGSGTVNLSEGDKLDLRDLLDHDGSRNQNDLKSLLSVFEDNEGGHLQVREGSSTSAITQEIVLMNHTFGTLTGGSATTSNQVIDYMLNNHMLDIDKP

**Corrected LapAj CDS:**

atgattaccaatgccattaccttggaacagggcgtaaccgtaacccaactcaaaggaaagatttacctgcttgccgcagacggcagccggaagctgttggttgaaggggatgtactgccaaaaggggctgtgattgtctctcccgatggtggcagctttatgggcggcaaccagtcgttcaatgtgcagccggccggtcagccgagcgagtcggccgaagagggagatgcaccccagctggcacaaaacggtgcacccggcactccgagcgatatcaatgcactgcaacaagccattttgggcggcgctgacccgaccaagactcttgaagctcctgcagcgggtggtgcacctgccgccggtggtggtggtggcgtcggcggtgttgctggcgccagcggtaacggcggctttgttactatcgaccgcgtaggggatgccactatctctcaggcggggtttgatacgacctacaataacggcaccgcgccgacgctggatgcgacaggcattgcagatcagcctttcatctccgccaatctgaccctctctgcgcagcctcaagtaaatgaaggcggttccatcacctttactgcctctgttgatgtacctgtttttggcaccgatctggtcattactctctcaaatggtgctgtcattaccattcctgctggccaaactaccggttccgttactgtggctgtgcctgccgacagcccctatgtcgacccgtccgcgatcactgtttcgatcactggtactcagggcggtggtttcaaccagatcattaccggtccttcaacaaccacccagattaacgacaccatcgataccaccacagtgaccctgggtgctccggcgcaggtcaacgaaggtgggcagatcacctacagcgccagtgtcaacaacgccccgcagagtgacctggtgctgaccctgtccaacggcgccagcatcaccatcaaggcgggcgagctgagcggcagcgtgacggtcgatgccccgagtgacgacgtttacaaagatggcagcagcctgacggtgagcatcaccggcagcagcggtggcaactacgagcagctggacaccagcagcaaggtgaccaccgaggtggccgacaccgtggacaccaccacggtgaccctgagcgcaccggcgcaggtcaacgaaggtgggcagatcacctacagcgccagtgtcaacaatgccccgcagagtgacctggtgctgaccctgtccaacggcgccagcatcaccatcaaggcgggcgagctgagcggcagcgtgacggtcgatgccccgagtgacgacgtttacaaagatggcagcagcctgacggtgagcatcaccggcagcagcggtggcaactacgagcagctggacaccagcagcaaggtgaccaccgaggtggccgacaccgtggacaccaccacggtgaccctgagcgcaccggcgcaggtcaacgaaggtgggcagatcacctacagcgccagtgtcaacaacgccccgcagagtgacctggtgctgaccctgtccaacggcgccagcatcaccatcaaggcgggcgagctgagcggcagcgtgacggtcgatgccccgagtgacgacgtttacaaagatggcagcagcctgacggtgagcatcaccggcagcagcggtggcaactacgagcagctggacaccagcagcaaggtgaccaccgaggtggccgacaccgtggacaccaccacggtgaccctgagcgcaccggcgcaggtcaacgaaggtgggcagatcacctacagcgccagtgtcaacaacgccccgcagagtgacctggtgctgaccctgtccaacggcgccagcatcaccatcaaggcgggcgagctgagcggcagcgtgacggtcgatgccccgagtgacgacgtttacaaagatggcagcagcctgacggtgagcatcaccggcagcagcggtggcaactacgagcagctggacaccagcagcaaggtgaccaccgaggtggccgacaccgtggacaccaccacggtgaccctgagcgcaccggcgcaggtcaacgaaggtgggcagatcacctacagcgccagtgtcaacaatgccccgcagagtgacctggtactgaccctgtccaacggcgccagcatcaccatcaaggcgggcgagctgagcggcagcgtgacggtcgatgccccgagtgacgacgtttacaaagatggcagcagcctgacggtgagcatcaccggcagcagcggtggcaactacgagcagctggacaccagcagcaaggtgaccaccgaggtggccgacaccgtggacaccaccacggtgaccctgagcgcaccggcgcaggtcaacgaaggtgggcagatcacctacagcgccagtgtcaacaatgccccgcagagtgacctggtgctgaccctgtccaacggcgccagcatcaccatcaaggcgggcgagctgagcggcagcgtgacggtcgatgccccgagtgacgacgtttacaaagatggcagcagcctgacggtgagcatcaccggcagcagcggtggcaactacgagcagctggacaccagcagcaaggtgaccaccgaggtggccgacaccgtggacaccaccacggtgaccctgagcgcaccggcgcaggtcaacgaaggtgggcagatcacctacagcgccagtgtcaacaacgccccgcagagtgacctggtgctgaccctgtccaacggcgccagcatcaccatcaaggcgggcgagctgagcggcagcgtgacggtcgatgccccgagtgacgacgtttacaaagatggcagcagcctgacggtgagcatcaccggcagcagcggtggcaactacgagcagctggacaccagcagcaaggtgaccaccgaggtggccgacaccgtggacaccaccacggtgaccctgagcgcaccggcgcaggtcaacgaaggtgggcagatcacctacagcgccagtgtcaacaatgccccgcagagtgacctggtactgaccctgtccaacggcgccagcatcaccatcaaggcgggcgagctgagcggcagcgtgacggtcgatgccccgagtgacgacgtttacaaagatggcagcagcctgacggtgagcatcaccggcagcagcggtggcaactacgagcagctggacaccagcagcaaggtgaccaccgaggtggccgacaccgtggacaccaccacggtgaccctgagcgcaccggcgcaggtcaacgaaggtgggcagatcacctacagcgccagtgtcaacaacgccccgcagagtgacctggtgctgaccctgtccaacggcgccagcatcaccatcaaggcgggcgagctgagcggcagcgtgacggtcgatgccccgagtgacgacgtttacaaagatggcagcagcctgacggtgagcatcaccggcagcagcggtggcaactacgagcagctggacaccagcagcaaggtgaccaccgaggtggccgacaccgtggacaccaccacggtgaccctgagcgcaccggcgcaggtcaacgaaggtgggcagatcacctacagcgccagtgtcaacaacgccccgcagagtgacctggtgctgaccctgtccaacggcgccagcatcaccatcaaggcgggcgagctgagcggcagcgtgacggtcgatgccccgagtgacgacgtttacaaagatggcagcagcctgacggtgagcatcaccggcagcagcggtggcaactacgagcagctggacaccagcagcaaggtgaccaccgaggtggccgacaccgtggacaccaccacggtgaccctgagcgcaccggcgcaggtcaacgaaggtgggcagatcacctacagcgccagtgtcaacaatgccccgcagagtgacctggtactgaccctgtccaacggcgccagcatcaccatcaaggcgggcgagctgagcggcagcgtgacggtcgatgccccgagtgacgacgtttacaaagatggcagcagcctgacggtgagcatcaccggcagcagcggtggcaactacgagcagctggacaccagcagcaaggtgaccaccgaggtggccgacaccgtggacaccaccacggtgaccctgagcgcaccggcgcaggtcaacgaaggtgggcagatcacctacagcgccagtgtcaacaatgccccgcagagtgacctggtgctgaccctgtccaacggcgccagcatcaccatcaaggcgggcgagctgagcggcagcgtgacggtcgatgccccgagtgacgacgtttacaaagatggcagcagcctgacggtgagcatcaccggcagcagcggtggcaactacgagcagctggacaccagcagcaaggtgaccaccgaggtggccgacaccgtggacaccaccacggtgaccctgagcgcaccggcgcaggtcaacgaaggtgggcagatcacctacagcgccagtgtcaacaacgccccgcagagtgacctggtgctgaccctgtccaacggcgccagcatcaccatcaaggcgggcgagctgagcggcagcgtgacggtcgatgccccgagtgacgacgtttacaaagatggcagcagcctgacggtgagcatcaccggcagcagcggtggcaactacgagcagctggacaccagcagcaaggtgaccaccgaggtggccgacaccgtggacaccaccacggtgaccctgagcgcaccggcgcaggtcaacgaaggtgggcagatcacctacagcgccagtgtcaacaacgccccgcagagtgacctggtgctgaccctgtccaacggcgccagcatcaccatcaaggcgggcgagctgagcggcagcgtgacggtcgatgccccgagtgacgacgtttacaaagatggcagcagcctgacggtgagcatcaccggcagcagcggtggcaactacgagcagctggacaccagcagcaaggtgaccaccgaggtggccgacaccgtggacaccaccacggtgaccctgagcgcaccggcgcaggtcaacgaaggtgggcagatcacctacagcgccagtgtcaacaacgccccgcagagtgacctggtgctgaccctgtccaacggcgccagcatcaccatcaaggcgggcgagctgagcggcagcgtgacggtcgatgccccgagtgacgacgtttacaaagatggcagcagcctgacggtgagcatcaccggcagcagcggtggcaactacgagcagctggacaccagcagcaaggtgaccaccgaggtggccgacaccgtggacaccaccacggtgaccctgagcgcaccggcgcaggtcaacgaaggtgggcagatcacctacagcgccagtgtcaacaatgccccgcagagtgacctggtactgaccctgtccaacggcgccagcatcaccatcaaggcgggcgagctgagcggcagcgtgacggtcgatgccccgagtgacgacgtttacaaagatggcagcagcctgacggtgagcatcaccggcagcagcggtggcaactacgagcagctggacaccagcagcaaggtgaccaccgaggtggccgacaccgtggacaccaccacggtgaccctgagcgcaccggcgcaggtcaacgaaggtgggcagatcacctacagcgccagtgtcaacaacgccccgcagagtgacctggtgctgaccctgtccaacggcgccagcatcaccatcaaggcgggcgagctgagcggcagcgtgacggtcgatgccccgagtgacgacgtttacaaagatggcagcagcctgacggtgagcatcaccggcagcagcggtggcaactacgagcagctggacaccagcagcaaggtgaccaccgaggtggccgacaccgtggacaccaccacggtgaccctgagctcggccaccaacggccagaccgtgaccgagggcggcagcattgtctacaccgccagcgtcagcaacccggtcaccggcagcccgctgaccgtgaccctctccaacggggtggtgatcaccattccggtgggcagcagcagcgctaacagtgacccggtaccggttcgtggagatgacatttatcagcaaggtgaagagttactgagcgtcactatcgacaagaccagtggtggcaattacgagaatgtggttagcactggcaccatcaccaatacagtggtggatgatcaggatttaccgactctgaccctgactggcgatgccaatgtggtcgaggggggcaaggcgagctacaccctgagcatcagcgatgcaccgaaaaccgatctgaccgtcaaggtggtggtgggccatatcaccaccgacgagggggatgtccaggcggtgacccgcgatgtggtcatcaaggcggggaccacctcggtgagcttcgacgtggcgaccaaggatgacgtctatgccgagccggccgagcagttcaaggtgagcgtgagcggcaccagtggcggtggctacgagaaggcgccggcgctgccgggtgcggtgaccaccaccatcactgacgaggatggcagccctgaccagccgaaggatgccccgaccctgaccctgaccggcgatgccaatgtggtcgaggggggcaaggcgagctacaccctgagcatcagcgatgcaccgaaaaccgatctgaccgtcaaggtggtggtgggccatatcaccaccgacgagggggatgtccaggcggtgacccgcgatgtggtcatcaaggcggggaccacctcggtgagcttcgacgtggcgaccaaggatgacgtctatgccgagccggccgagcagttccgggtgagcgtgagcggcaccagtggcggtggctacgagaaggcgccggcgctgccgggtgcggtgaccaccaccatcactgacgaggatggcagccctgaccagccgaaggatgccccgaccctgaccctgaccggcgatgccaatgtggtcgaggggggcaaggcgagctacaccctgagcatcagcgatgcaccgaaaaccgatctgaccgtcaaggtggtggtgggccatatcaccaccgacgagggggatgtccaggcggtgacccgcgatgtggtcatcaaggcggggaccacctcggtgagcttcgacgtggcgaccaaggatgacgtctatgccgagccggccgagcagttccgggtgagcgtgagcggcaccagtggcggtggctacgagaaggcgccggcgctgccgggtgcggtgaccaccaccatcactgacgaggatggcagccctgaccagccgaaggatgccccgaccctgaccctgaccggcgatgccaatgtggttgaggggggcaaggcgagctacaccctgagcatcagcgatgcaccgaaaaccgatctgaccgtcaaggtggtggtgggccatatcaccaccgacgagggggatgtccaggcggtgacccgcgatgtggtcatcaaggcggggaccacctcggtgagcttcgacgtggcgaccaaggatgacgtctatgccgagccggccgagcagttccgggtgagcgtgagcggcaccagtggcggtggctacgagaaggcgccggcgctgccgggtgcggtgaccaccaccatcactgacgaggatggcagccctgaccagccgaaggatgccccgaccctgaccctgaccggcgatgccaatgtggttgaggggggcaaggcgagctacaccctgagcatcagcgatgcaccgaaaaccgatctgaccgtcaaggtggtggtgggccatatcaccaccgacgagggggatgtccaggcggtgacccgcgatgtggtcatcaaggcggggaccacctcggtgagcttcgacgtggcgaccaaggatgacgtctatgccgagccggccgagcagttcaaggtgagcgtgagcggcaccagtggcggtggctacgagaaggcgccggcgctgccgggtgcggtgaccaccaccatcactgacgaggatggcagccctgaccagccgaaggatgccccgaccctgaccctgaccggcgatgccaatgtggtcgaggggggcaaggcgagctacaccctgagcatcagcgatgcaccgaaaaccgatctgaccgtcaaggtggtggtgggccatatcaccaccgacgagggggatgtccaggcggtgacccgcgatgtggtcatcaaggcggggaccacctcggtgagcttcgacgtggcgaccaaggatgacgtctatgccgagccggccgagcagttccgggtgagcgtgagcggcaccagtggcggtggctacgagaaggcgccggcgctgccgggtgcggtgaccaccaccatcactgacgaggatggcagccctgaccagccgaaggatgccccgaccctgaccctgaccggcgatgccaatgtggttgaggggggcaaggcgagctacaccctgagcatcagcgatgcaccgaaaaccgatctgaccgtcaaggtggtggtgggccatatcaccaccgacgagggggatgtccaggcggtgacccgcgatgtggtcatcaaggcgggcaccacctcggtgagcttcgacgtggcgaccaaggatgacgtctatgccgagccggccgagcagttccgggtgagcgtgagcggcaccagtggcggtggctacgagaaggcgccggcgctgccgggtgcggtgaccaccaccatcactgacgaggatggcagccctgaccagccgaaggatgccccgaccctgaccctgaccggcgatgccaatgtggttgaggggggcaaggcgagctacaccctgagcatcagcgatgcaccgaaaaccgatctgaccgtcaaggtggtggtgggccatatcaccaccgacgagggggatgtccaggcggtgacccgcgatgtggtcatcaaggcgggcaccacctcggtgagcttcgacgtggcgaccaaggatgacgtctatgccgagccggccgagcagttccgggtgagcgtgagcggcaccagtggcggtggctacgagaaggcgccggcgctgccgggtgcggtgaccaccaccatcactgacgaggatggcagccctgaccagccgaaggatgccccgaccctgaccctgaccggcgatgccaatgtggttgaggggggcaaggcgagctacaccctgagcatcagcgatgcaccgaaaaccgatctgaccgtcaaggtggtggtgggccatatcaccaccgacgagggggatgtccaggcggtgacccgcgatgtggtcatcaaggcgggcaccacctcggtgagcttcgacgtggcgaccaaggatgacgtctatgccgagccggccgagcagttcaaggtgagcgtgagcggcaccagtggcggtggctacgagaaggcgccggcgctgccgggtgcggtgaccaccaccatcactgacgaggatggcagccctgaccagccgaaggatgccccgaccctgaccctgaccggcgatgccaatgtggtcgaggggggcaaggcgagctacaccctgagcatcagcgatgcaccgaaaaccgatctgaccgtcaaggtggtggtgggccatatcaccaccgacgagggggatgtccaggcggtgacccgcgatgtggtcatcaaggcggggaccacctcggtgagcttcgacgtggcgaccaaggatgacgtctatgccgagccggccgagcagttccgggtgagcgtgagcggcaccagtggcggtggctacgagaaggcgccggcgctgccgggtgcggtgaccaccaccatcactgacgaggatggcagccctgaccagccgaaggatgccccgaccctgaccctgaccggcgatgccaatgtggtcgaggggggcaaggcgagctacaccctgagcatcagcgatgcaccgaaaaccgatctgaccgtcaaggtggtggtgggccatatcaccaccgacgagggggatgtccaggcggtgacccgagatgtggtcatcaaggcggggaccacctcggtgagcttcgacgtggcgaccaaggatgacgtctatgccgagccggccgagcagttcaaggtgagcgtgagcggcaccagtggcggtggctacgagaaggcgccggcgctgccgggtgcggtgaccaccaccatcactgacgaggcggtaccgggagcggaagacaccgtgtatgcccagatcgaggtggatcaatccagcgtggtggaaggtggcgcgctcaaatacaccgtgagcctggtggacaaggacggcaatgccgtgacactgccggcaggcaaagtggtgaccctgaacttggcatggagcggggcggcggccaatgatggcgacaccagcggtcgtccggccagcgtgaccatcggtgcaaacggcaaagccgaatttactgttactacagtaaacgacaccatttatgagatggatgaaaagcttgctgtcactatcagtggtgtgaagatcaattcagcatttgaagctatttcgatagtaaatggaaaaggtagtgctgagaccgtcattgtggacaacgacgctccccctaccgtggtgatcaataaacctgcggtggtgtccgaagaggggttggccaacgggttgcctgataacgcaggtaatacagataccaccaatgcgaccgtcgccagcggcactattaccgtgggggatgtcgatagcagtagcgtgacggtcagcctctctggtcgatccggactcaagtccggcgggctggatgtgaagtggaactgggatgcatcaggtaaagtgttgaccgggtataccggtaccttgggtggtgcggactataaggctgtgcttgatgtcaaactgaccgcgcctgcaggaagtactaaaggagattggggctatgacgttactctgaaaggagctctcgatcatccggtgactggtgccgaggacacgctgagcttcaacgtcggagtttctgtttctgatggacagagcactacggtcggttcgctgcctatcacagttgaggatgacgcgccaatcgctggtgacatggcggcagtatctctggtgaaaaccagtatcccggatgtgttgaccggcaagttcagcctgaccgggtatgtcggggatagcggcagcattgatgcgggcaaattcactatcacggcacggggattcgagtcttcgacttcctccaggctcgttgatgccaaggtcaatgcgtccgttgacgggctaggcgtcaagagcacgagcgcgccttatcacaacattgataatgagatcgatttccgcaaatttgctgatggttcgagtgtatccgaggagttggtgatcaagctggatgcgggtactgtggcatatggcgccaagatcgagttcagcaagatctatggtgcagaactcgagtgcggtgtggtcgagttctggcgcgacggtaaactggttgcaacacagacattcagctcgaatgctgccaatggtgattatgctgccaacttccaggttcagcagggcggttttgacaccatggtgatcaaggcgaccgataacggcaagtcggccagctggggtgacaacagtgactttactgtcaagtctatcgagttccttggttcctctaccccgcaggcgattgcctatggttcgggcacgctcgatccgcagtggggggctgatggtaaaggacgtttcgaactggtcaccggtacggtcgaatccggcctcaagacagttgccggtctggctatcgcgatcaccgccgatggggctaataccctgcttggtaaagacagcaatggcagtctgatctttaagatggagttcacacctgccacaggcaaatgggagttcttccagtatgccgagatgcagcgtcctgctgacggtgatattgactttacatttaaagcttacgatcgcgatggtgatggctcgcaaggcagtttcgcggtcaatccgctggttcgccccgatgtaatgggggtgtcgaatgccagtgtgactgaggggggggctctgcaacatgttgtcaccctgagcggggcgaccaatgtggctaccgagtacagcctctccattgctggcagcggtgctcatgctgccagcagtgctgattggggagctttgcagtttaccaacggggtgacctacaacagcaccaccggcaagataaccgtgcctgccggagtatcgagtttcactgtatcgctgaatacggtaaacgacacactgcccgagttcacagaaacgctggcgatctcggtagggggcgtttccggtaccggcaccatagtcgacaatgatctctcggtgcggctcggtgtcggtctggttgatgaggatggtttgtctggtggcaacgccggtttgcccgagcctgtcggggcgccgaccagcggtccgctctcggttacccaatcgctgcaggttaccgatggcaatggtagcaacgtatccggcgttcagctcaagctggtcagcattaccggcttgagcggcattaccggcattgatggccagcccgtgaatgtggtgcaggatggtgcgggtctcaaaggttactttggaaccaacccggccaatgtggcctttaccgtaacggtagacaacaattcgaatccaccgagctataccttcaccctgatcaagccgctcagccaccttgtggatggtcaaagcagcgtattgagcagtcaggatgagctgcgtctcacggtgaattatgaggtgagcaaagctggttccgacactgcgagcggctctttcgatgtggccgtgcgtgatgatgtgccggttgctgtggtcgatgatgccacgcttaacgtggtggtagattcgttccagttctccggtgttgaggcgtcctggatcaatgttgttggtggtaagaatctggagtatttcgatggtccggataacgactcggccaatgatcagatccgctggggtggcaagaatgaagttaagtcaggttacggattcgctgacaacgatgctgcgctgaatgggcagatcccgctcaacgaagagatcaagttgggtacctttacccactacaactatccgatctcttccggtacatcgattagtgctgtcaccatgaaggcgatattctcggtcaccgattccatgggaagagtgactcccgtgactgtcaccatcaactttaatcacaacgaaacgccaaacgatggggccgatccgcgagatatcatcactatcggaactgcgaccgcgagcttcaactttgaaggcaaggtctacagtcttgaggtgctgggcttccgcgataccggtggcaacgtagtgaagactatctacaccaacgaaaatgccagcaacagcttcgatctgatcgtcaagttggtagagggttctggttatcagttgccacagactacaggcaatgtgctggccaacgatcttgccggtgctgatggtggtttgagtgtcattggctacggagtaggcagcagctcgaccacctataccaatggtgcgggtacccaggtagttggtctgtacggggtgctgaccattctggccaacggtgcctatacctatcaggtcaccaagaatggcagccaaatcccggctgatgcacgcgaggtgttcagctacagcgtgcgggatggcgatggcgacactaccaacagtactctgaccatcagcgtcaatccggtggacagcaacggggtaccggttcatctgccgttgacggtagatggtaccgacctcaacgactctatcgttgtgcgcaatggcgaaaatgcggctaaccctgatcgcctggatgttgcctttggtgggaatctgctgggcgaggtcaccaccagtaacggcggtacagaccatgtgcataccggtatcagttacaacaaagggagcctcgatcaggtcgtcagtggtggtgcaggcaatgaccacatcgaaaccggcagcggcaacgatgtgatctatgcaggcaaaaccggggcggatggatttggtaccgacgattcgttgcaactgacggtagctcagctcaaggctcatcacatcatgacaggcagcttgtctggcactgatgccatgctcgatggcgacggtctgttgctgagtgttgatgtcagcagtagcaaagcagatgtggtcaatggcggtagcggtaatgacaagatctacggacaatcaggttcggatatcttgtttggtggtaccggtgacgactatatcgatggcggcagtcacaacgatgcactgcgtggcggtctggggaacgatatcctgatcggtgggctcggcaacgatgtgatgcgtggtgacggcggctcagatactttcgtctggaatcagggggatactgtatccggttctctgaccaaggattacatcatggacttcaacaaggggtcaggcacagtcaatctttcggaaggtgacaaactggatctgcgcgatctgctggatcacgacggttctcgcaaccagaacgatctgaagagcctgctctcggtgtttgaggacaatgaaggggggcatctgcaggtcagggagggcagctctacttctgcgattacccaagaaatcgtgttgatgaaccacacctttggcacgctgacgggcggttcggcgaccacgtccaatcaggtgatcgattacatgttgaataatcacatgctcgatattgacaagccataa
