## Supplementary Information (Figs, Tables, and References) for "*Aeromonas* adhesins facilitate kin and non-kin attachment to enable T6SS-mediated antagonism in liquid"

**Supplementary Figure S1-S5**

**Supplementary Table S1-S2**

**Supplementary Files S1-S2 (captions)**

**Supplementary References**

SUPPLEMENTARY FIGURES

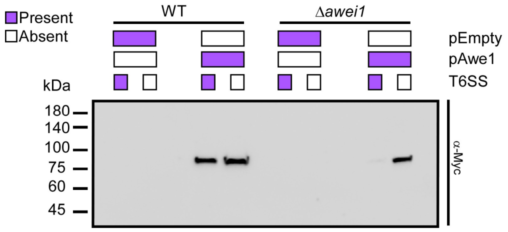


**Figure S1. Inducible expression of Awe1 from a plasmid in *Aj* strains.** The expression of a C-terminally Myc-tagged Awe1 from an arabinose-inducible plasmid in the indicated strains used in Fig. 1A.

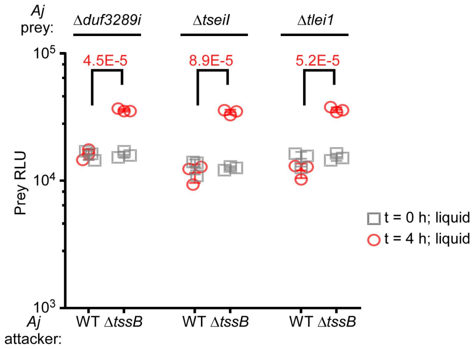


**Figure S2. *Aeromonas jandaei* deploys all known T6SS effectors during T6SS-mediated prey intoxication in liquid media.** LiQuoR-based viability measurements (RLU) of *Aj* prey strains lacking the indicated T6SS effector and immunity pair before (t = 0 h) and after (t = 4 h) co-incubation with the indicated *Aj* attacker at a 4:1 (attacker:prey) ratio in LB media at 30˚C. Data are shown as the mean ± SD; n = 3 samples. The statistical significance between samples at the 4 h time point (for each prey strain) was calculated using an unpaired, two-tailed Student's *t* test. The experiment was repeated three times with similar results. Results from a representative experiment are shown.

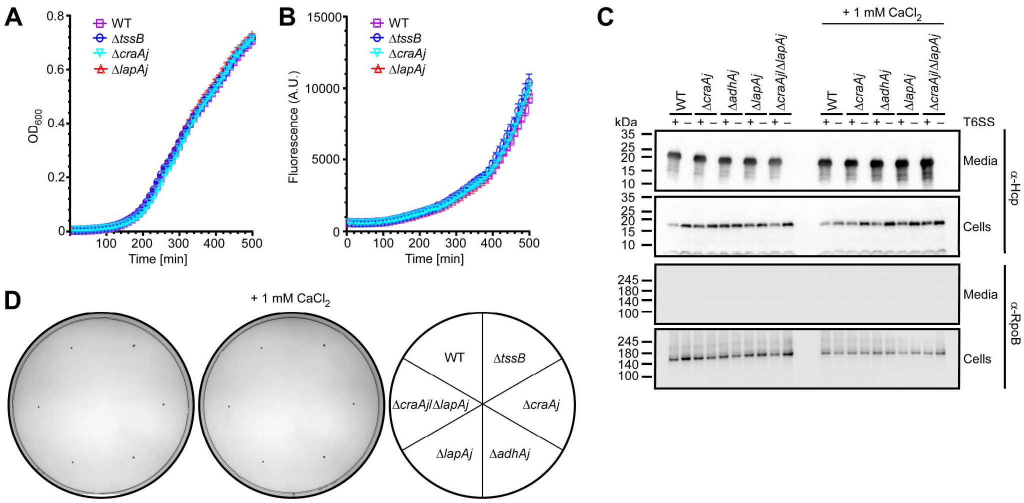


**Figure S3. Adhesin deletions do not affect *Aj* growth, T6SS activity, or swimming. (A-B)** Growth of the indicated *Aj* strains containing a plasmid for the constitutive expression of the red fluorescent protein mLychee in LB media at 30˚C, as measured by optical density at 600 nm ($OD_{600}$) readings (A) or red fluorescence (arbitrary units; A.U.) readings (B). Data are shown as the mean ± SD; n = 3 samples. **(C)** Expression (cells) and secretion (media) of Hcp from wild-type (WT; $T6SS^+$) or Δ*tssB* ($T6SS^-$) *Aj* strains grown for 3 h in LB media at 30ºC, with or without 1 mM $CalCl_2$. RNA polymerase beta subunit (RpoB) was used as a loading and lysis control. **(D)** Motility of the indicated *Aj* strains in soft agar (0.3% [w/v]) swimming plate. The experiments were repeated three times with similar results. Results from a representative experiment are shown.

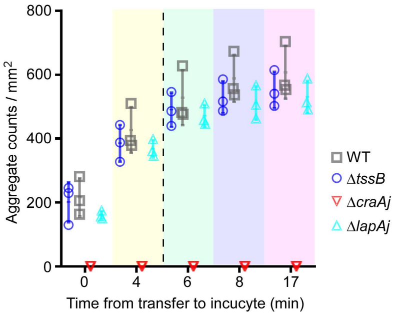


**Figure S4. Accumulation of *Aeromonas* auto-aggregates at the bottom of a 96-well plate peaks within five minutes.** Quantification of *Aj* aggregates from microscope images at the indicated time points after the 96-well plate was placed in the Incucyte, following a 1-hour incubation with shaking, as described in Figure 4A-C. The vertical dashed line denotes the 5-minute timepoint used to acquire images in Figure 4B-C.

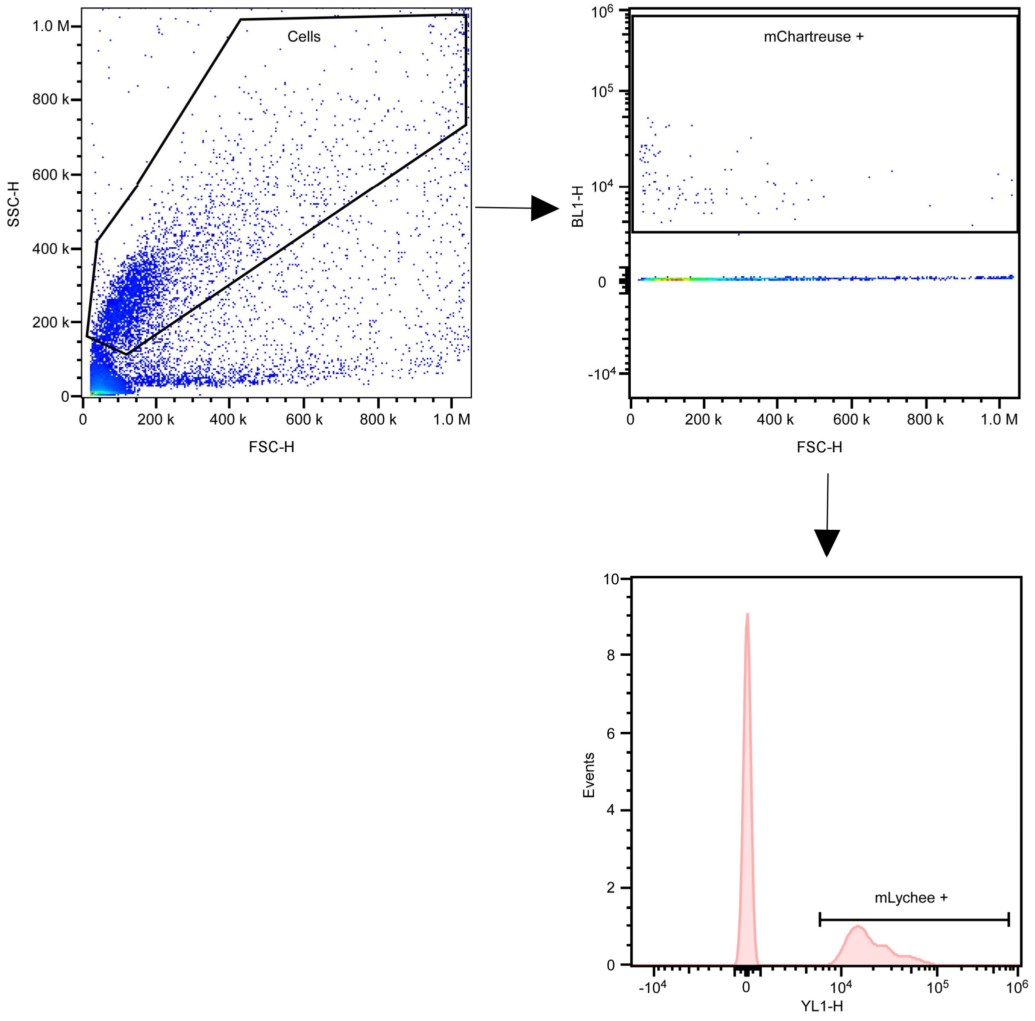


**Figure S5. *Aj* and *E. coli* co-aggregation gating strategy.** Representative flow cytometry plots showing the gating strategy to isolate the co-aggregates (red-positive events within the green-positive population).

SUPPLEMENTARY TABLES

**Table S1: Bacterial strains used in this study.**

| Strain Name | Genotype | Comments | Source |
|---|---|---|---|
| *Aeromonas jandaei* DSM 7311 | Wild-type | Used for all assays in this study and for generating deletion strains | DSMZ collection |
| *Aj* Δ*tssB* | DSM 7311 Δ*we862_rs13035* | Used as a T6SS⁻ strain in growth, competition, expression, secretion, and auto-aggregation assays | [1] |
| *Aj* Δ*awei1* | DSM 7311 Δ*we862_rs20675–we862_rs20665* | Deletion of the *awe1* effector and its cognate immunity genes. Used as prey in competition assays | [2] |
| *Aj* Δ*tssB*/Δ*awei1* | DSM 7311 Δ*we862_rs13035* / Δ*we862_rs20675–we862_rs20665* | Used in expression and growth assays | This study |
| *Aj* Δ*tseiI* | DSM 7311 Δ*we862_rs13125–we862_rs13130* | Deletion of the *tseI* effector and its cognate immunity gene. Used as prey in competition assays | [2] |
| *Aj* Δ*tlei1* | DSM 7311 Δ*we862_rs16925–we862_rs16920* | Deletion of the *tlei1* effector and its cognate immunity gene. Used as prey in competition assays | [2] |
| *Aj* Δ*duf3289i* | DSM 7311 Δ*we862_rs09995* + downstream immunity (not annotated on NCBI) | Deletion of the *duf3289* effector and its cognate immunity gene. Used as prey in competition assays | [2] |
| *Aj* Δ*craAj* | DSM 7311 Δ*we862_rs13595* | Used in competition, secretion, growth, swimming, and auto-aggregation assays | This study |
| *Aj* Δ*tssB*/Δ*craAj* | DSM 7311 Δ*we862_rs13035* / Δ*we862_rs13595* | Used in competition, secretion, and co-aggregation assays | This study |
| *Aj* Δ*craAj*/Δ*awei1* | DSM 7311 Δ*we862_rs13595* / Δ*we862_rs20675–we862_rs20665* | Used in secretion assays and as prey in competition assays | This study |

| | | | |
|---|---|---|---|
| *Aj* Δ*adhAj* | DSM 7311 Δ*we862_rs16715* | Used in competition and secretion assays | This study |
| *Aj* Δ*tssB*/Δ*adhAj* | DSM 7311 Δ*we862_rs13035* / Δ*we862_rs16715* | Used in competition and secretion assays | This study |
| *Aj* Δ*lapAj* | DSM 7311 Δ*we862_rs21075* | Used in competition, secretion, growth, swimming, and auto-aggregation assays | This study |
| *Aj* Δ*tssB*/Δ*lapAj* | DSM 7311 Δ*we862_rs13035* / Δ*we862_rs21075* | Used in competition and secretion assays | This study |
| *Aj* Δ*craAj*/Δ*adhAj* | DSM 7311 Δ*we862_rs13035* / Δ*we862_rs16715* | Used in competition assays | This study |
| *Aj* Δ*tssB*/Δ*craAj*/Δ*adhAj* | DSM 7311 Δ*we862_rs13035* / Δ*we862_rs13595* / Δ*we862_rs16715* | Used in competition assays | This study |
| *Aj* Δ*craAj*/Δ*lapAj* | DSM 7311 Δ*we862_rs13595* / Δ*we862_rs21075* | Used in competition and swimming assays | This study |
| *Aj* Δ*tssB*/Δ*craAj*/Δ*lapAj* | DSM 7311 Δ*we862_rs13035* / Δ*we862_rs13595* / Δ*we862_rs21075* | Used in competition and co-aggregation assays | This study |
| *Aj* Δ*adhAj*/Δ*lapAj* | DSM 7311 Δ*we862_rs16715* / Δ*we862_rs21075* | Used in competition assays | This study |
| *Aj* Δ*tssB*/Δ*adhAj*/Δ*lapAj* | DSM 7311 Δ*we862_rs13035* / Δ*we862_rs16715* / Δ*we862_rs21075* | Used in competition assays | This study |
| *Vibrio parahaemolyticus* RIMD 2210633 | Wild-type | Used in competition assays | Obtained from Kim Orth |
| *Vibrio parahaemolyticus* Δ*hcp1*/Δ*hcp2* | RIMD 2210633 Δ*vp1393*/Δ*vpa1027* | Used in competition assays | [3] |
| *Vibrio coralliilyticus* ATCC BAA-450 | Wild-type | Used in competition assays | ATCC collection |
| *Vibrio coralliilyticus* ATCC BAA-450 Δ*hcp1*/Δ*tssM2* | ATCC BAA-450 Δ*VIC_RS16330*/Δ*vic_RS20055* | Used in competition assays | [4] |

| | | | |
|---|---|---|---|
| *Escherichia coli* DH5α (λ-pir) | K-12 derivative laboratory strain containing λ pir | Used for plasmid construction and maintenance, and as prey in competition assays | Obtained from Eric V. Stabb |
| *Escherichia coli* HB101 | K-12 derivative laboratory strain | Harbors pRK2013; Used as a conjugation helper strain | ATCC collection |

### Table S2: Plasmids used in this study.

| Plasmid name | Description | Purpose | Source |
|---|---|---|---|
| pDM4 | $Cm^R$, suicide vector with $oriV_{R6K}$ | Used as a backbone when constructing plasmids for gene deletions | 5 |
| pDM4:*awiU-awe1-awiD* | pDM4 containing 1 kb upstream of the gene encoding AwiU and 1 kb downstream of the gene encoding AwiD in *Aj* in its MCS | Used to construct *awei1* deletions in *Aj* strains | 2 |
| pDM4:*craAj* | pDM4 containing 1 kb upstream and 1 kb downstream of the gene encoding CraAj in *Aj* in its MCS | Used to delete *craAj* in *Aj* strains | This study |
| pDM4:*adhAj* | pDM4 containing 1 kb upstream and 1 kb downstream of the gene encoding AdhAj in *Aj* in its MCS | Used to delete *adhAj* in *Aj* strains | This study |
| pDM4:*lapAj* | pDM4 containing 1 kb upstream and 1 kb downstream of the gene encoding LapAj in *Aj* in its MCS | Used to delete *lapAj* in *Aj* strains | This study |
| $pBAD^K$/*Myc*-His | pBR322 ori-containing plasmid harboring a $Kan^R$ cassette, *araC*, and an MCS following a P*bad* promoter. A Myc-His tag is encoded at the 5' end of the MCS. | Used as an empty plasmid control for growth and expression assays in *Aj* | 6 |
| pAwe1 | $pBAD^K$/*Myc*-His plasmid containing *awe1*, in-frame with a C-terminal Myc-His tag | Used for arabinose-inducible expression of Awe1 | 2 |
| pAKlux2 | pBBR1MCS4-based plasmid expressing luxCDABE from the lacZ promoter (constitutive in the absence of LacI) | Used as a template for constructing pCmLux2 | Addgene collection 7 |
| $pBAD33.1^F$ | pBAD33.1 with a FLAG tag inserted at the 3' end of the MCS | Used as a template for amplifying the chloramphenicol resistance cassette to construct pCmLux2 | 8 |

| | | | |
|---|---|---|---|
| pCmLux2 | pAKlux2 derivative in which the ampicillin resistance cassette was replaced with a chloramphenicol resistance cassette from pBAD33.1[F] | Used to monitor prey luminescence in competition assays | This study |
| pNF02-mLychee | Enhanced red fluorescent protein, derived from mApple, under the constitutive promoter proDp | Used as a template for amplifying mLychee and proDp for constructing pBAD33.4:mLychee | Addgene collection [9] |
| pNF02-mChartreuse | Enhanced green fluorescent protein, derived from sfGFP, under the constitutive promoter proDp | Used as a template for amplifying mChartreuse and proDp for constructing pBAD33.4:mChartreuse | Addgene collection [9] |
| pBAD33.1 | A mobilizable plasmid for arabinose-inducible protein expression, with ori15A and chloramphenicol resistance | Used as a template for constructing the pBAD33.4 plasmid | Addgene |
| pBAD33.4 | pBAD33.1-based plasmid in which the arabinose-inducible promoter was replaced by the constitutive proDp promoter; a AAAGGG linker and a C-terminal FLAG tag were inserted at the 3′ end of the MCS | Used for constitutive protein expression | This study |
| pBAD33.4:mLychee | pBAD33.4 plasmid containing the CDS of mLychee from pNF02-mLychee, in-frame with a AAAGGG linker and a FLAG tag at the C-terminus | Used for constitutive mLychee expression in *A j* strains | This study |
| pBAD33.4:mChartreuse | pBAD33.4 plasmid containing the CDS of mChartreuse from pNF02-mChartreuse, in-frame with a AAAGGG linker and a FLAG tag at the C-terminus | Used for constitutive mChartreuse expression in *E. coli* | This study |

SUPPLEMENTARY FILES (captions)

**File S1. Sanger sequence of the region containing an erroneous insertion in *lapAj*.**

**File S2. Complete amino acid sequence of LapAj.**
